# Delivery of small interfering RNA and antisense oligonucleotides across the blood-brain barrier with monovalent transferrin receptor 1 binding VHH-Fc fusion proteins

**DOI:** 10.64898/2026.08.13.744307

**Authors:** Ian J. Huggins, Michele Carrer, Jinro A. Santos, Michael Fazio, Bret Holguin, Stevee Phi, Thazha P. Prakash, Megan Afetian, Mohsen A. Bakooshli, Stephanie K. Klein, Rodrigo Galindo-Murillo, Andrew A. Rodriguez, Fredrik Kamme, Hans Gaus, Alfred Chappell, Mariana Bravo-Hernandez, Antonio Pinto-Duarte, Ryan Quinones, Guillaume Jacquot, Marion David, Frank Rigo, Holly B. Kordasiewicz, Hien T. Zhao, Paymaan Jafar-nejad, Michael Tanowitz, Eric E. Swayze

**Affiliations:** Ionis Pharmaceuticals, 2855 Gazelle Court, Carlsbad, CA 92010, United States; VECT-HORUS, Faculté de la Timone, 27 Boulevard Jean Moulin, 1er étage aile Verte, 13005 Marseille, France

**Keywords:** Blood-brain-barrier, Transferrin receptor, Receptor-mediated transcytosis, VHH, siRNA, ASO

## Abstract

The blood-brain barrier (BBB) is a highly selective cell layer that restricts the diffusion of diverse chemical entities into the central nervous system (CNS) from systemic circulation. Macromolecular therapeutics including oligonucleotides, peptides, and monoclonal antibodies exhibit only minimal brain distribution after systemic dosing due to exclusion by the BBB. Receptor-mediated transcytosis (RMT) has evolved to transport vital cargo across the BBB through a specialized vesicular transport pathway. Transferrin receptor 1 (TfR1) shuttles transferrin, its natural ligand, across the BBB, as well as TfR1-binding IgG antibodies and conjugates. Here, we describe a novel monovalent TfR1-binding VHH-Fc for the delivery of oligonucleotide cargo, including antisense oligonucleotides (ASOs) and small interfering RNAs (siRNAs) across the BBB in rodents and non-human primates (NHPs), supporting the translational potential of the VHH-antisense RMT platform for the treatment of neurological disorders. We explore the role of binding affinity, conjugation site, drug-antibody ratio (DAR), and conjugation chemistry, and determine that binding affinity, DAR and conjugation site are major determinants of RMT capacity and brain activity of siRNAs delivered across the BBB.

**Graphical Abstract / Highlights:** 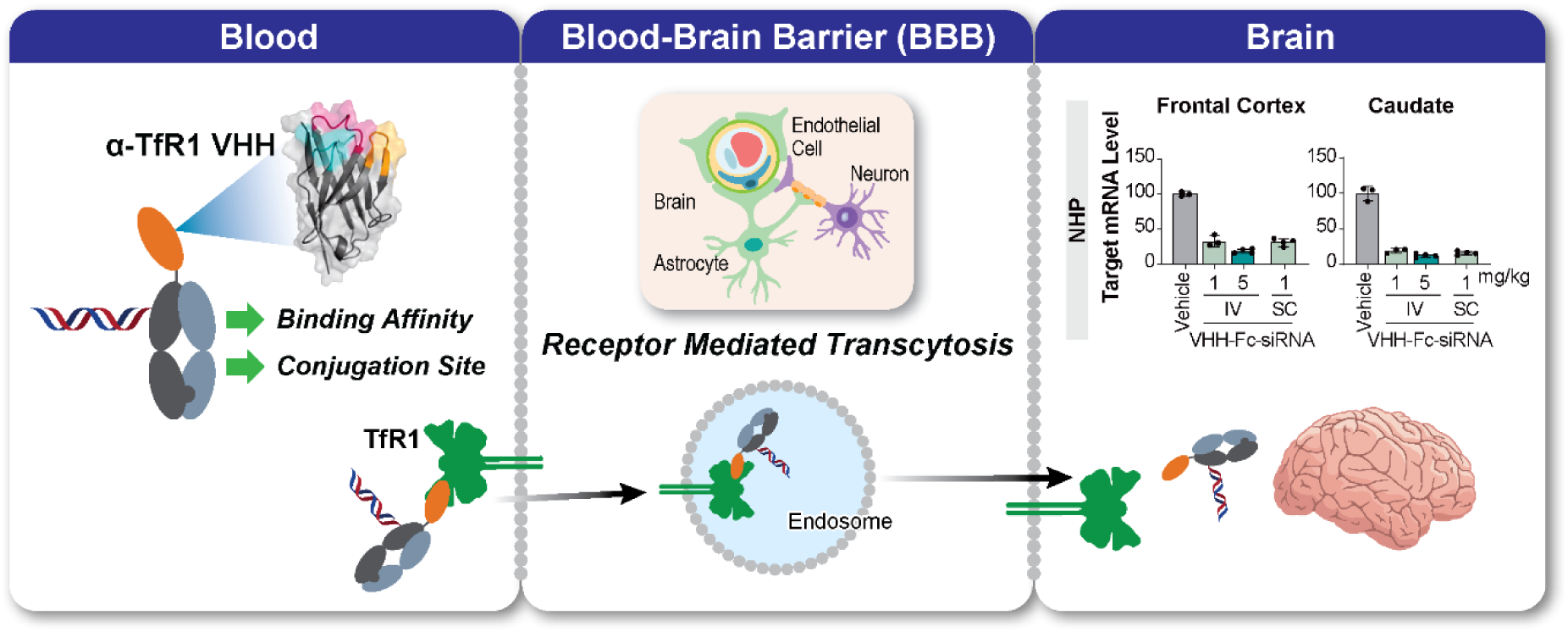

‒ Anti-TfR1 (α-TfR1) VHH ligands formatted as heterodimeric, 2-chain monovalent VHH-Fc were engineered for conjugation to siRNA and ASO.
‒ Systematic in vivo evaluation of VHH clones spanning a range of TfR1 binding affinities revealed a relationship between TfR1 binding affinity and the CNS activity of intravenously dosed VHH-Fc-siRNA conjugates.
‒ By optimizing TfR1 binding affinity, conjugation site, and conjugation chemistry, we identified VHH-Fc-siRNA molecules that efficiently cross the BBB via receptor-mediated transcytosis and reduce target mRNA across CNS tissues, including deeper brain regions, after intravenous (IV) or subcutaneous (SC) dosing in mice and non-human primates (NHPs).

## INTRODUCTION

Oligonucleotide therapeutics, including antisense oligonucleotides (ASOs) and small interfering RNAs (siRNAs) enable potent, selective and durable modulation of gene expression for the treatment of disease. Chemical conjugation to tissue targeting ligands, including GalNAc (1, 2), GLP-1 peptides (3, 4), cyclic peptides (5), lipids (6, 7), monoclonal antibodies (8) and antibody fragments (9) has been utilized to improve the pharmacokinetic and pharmacodynamic profiles of these drugs in multiple tissues, including liver, pancreatic beta cells, skeletal and cardiac muscle, adipose tissue, and brain following administration by intravenous (IV) or subcutaneous (SC) routes.

Tissues of the central nervous system (CNS) are protected by the blood-brain barrier (BBB), a layer of specialized cells, including endothelial cells connected by tight junctions that prevent passive diffusion of macromolecules such as oligonucleotides, many small molecules, and ions into the CNS from systemic circulation (10). The BBB employs receptor mediated transcytosis (RMT) to carry out selective transport of vital nutrients, growth factors, and other important molecules from the blood to the brain (11, 12). Transferrin receptor 1 (TfR1), encoded by the TFRC gene, is a rapidly recycling endocytic receptor that functions to carry iron-loaded holo-transferrin (Tf) from circulation into cells (13, 14) and is abundantly expressed on the surface of the endothelial cells of the BBB (15–17). Normally, following endocytosis, iron is released into the cell and the TfR1-Tf complex is recycled to the cell surface, where iron-free apo-Tf is released into circulation (18). In the endothelial cells of the BBB, the TfR1-Tf complex enters the transcytosis pathway, and Tf is released intact across the BBB via RMT (14). Other molecules that bind TfR1, including monoclonal antibodies, can also accumulate in the brain via RMT (19).

The inability of ASOs and siRNAs to efficiently traverse the BBB necessitates direct delivery to the cerebrospinal fluid, most commonly via intrathecal (IT) injection, for the treatment of neurological disorders. IT administration supports widespread distribution, cellular uptake, and target engagement throughout the CNS in preclinical models (20, 21) and humans (22, 23), and is employed to deliver clinically validated disease modifying therapeutics including nusinersen and tofersen (24–27). Although IT administration is clinically validated and can achieve broad CNS exposure, it remains an invasive procedure and may be less practical for chronic repeat dosing, particularly in large patient populations or in diseases requiring long-term treatment. A BBB-penetrant RNA therapeutic platform could enable systemic administration, improve delivery to brain regions that are less accessible from the cerebrospinal fluid, and expand the therapeutic potential of oligonucleotide medicines.

Chemical conjugation of ASOs to TfR1-binding IgG ligands has been shown to improve tissue pharmacokinetics (PK) in deep brain regions by enabling these molecules to cross the BBB via RMT (28). Naturally derived anti-TfR1 IgG antibodies comprise 2 identical heavy chains (HC) and 2 identical light chains (LC) that assemble via non-covalent association and interchain disulfide bonds, forming Y-shaped molecules with distinct functional domains (29–31). Two identical fragment antigen binding (Fab) arms contain the variable complementarity determining regions (CDRs) of the heavy and light chains that mediate the unique molecular target recognition of individual antibody clones. Fab arms connect via a flexible hinge to the fragment crystallizable (Fc) region, a constant domain that mediates immune effector functions through Fc-gamma receptor (FcγR) interactions and long circulation half-life through neonatal Fc receptor (FcRn) interactions (32). Natural anti-TfR1 IgG antibodies are thus bivalent and capable of binding and clustering receptors, which hampers RMT and increases lysosomal degradation of TfR1 (33). Engineered monovalent anti-TfR1 IgG antibodies bind the receptor with lower apparent affinity than their parent bivalent antibodies, do not induce receptor clustering, and more efficiently traverse the BBB in rodent and non-human primate (NHP) models (34–36). However, these molecules are comparatively complex to produce, typically requiring assembly of heterodimeric Fc regions in which one heavy chain lacks a Fab arm, while the other retains a Fab that must be co-expressed and correctly assembled with a cognate light chain. Eliminating the requirement for the additional light chain would substantially simplify the design of monovalent antibody-based BBB ligands, improve manufacturability, and reduce the total dose of antibody–oligonucleotide conjugates required to achieve therapeutic benefit.

Camelids, including camels, llamas, alpacas and other related species, produce HC and LC containing IgG antibodies, but also produce a class of HC-only antibody (HCAb), which mediate antigen binding without light chains (37, 38). The antigen binding region is referred to as the variable heavy domain of HCAbs (VHH) (39). VHH domains are capable of picomolar binding affinities and can be recombinantly expressed as single-domain antibodies (sdAbs). VHHs possess several key features differentiating them from traditional Fab or single-chain fragment variable (scFv) antibody formats, including low molecular weight (approximately 15 kDa), a more compact binding surface that may recognize epitopes inaccessible to monoclonal antibodies (mAbs), and simple production, which enables rapid structure-activity relationship (SAR) cycles, and scalable manufacturing. In this work, we employ anti-TfR1 VHH ligands formatted as heterodimeric, 2-chain monovalent VHH-Fc of approximately 65 kDa molecular weight to deliver siRNAs and ASOs across the BBB in vivo. Via optimization of TfR1 binding affinity, conjugation site, and conjugation chemistry, we observe widespread target reduction in CNS tissues, including deeper brain regions, following IV or SC dosing of VHH-Fc-siRNA and VHH-Fc-ASO in mice and NHPs (Graphical abstract).

## MATERIALS AND METHODS

### Solid phase synthesis of amino modified ASOs and siRNAs

All oligonucleotides were synthesized on an ÄKTA Oligopilot™ synthesizer 10 at 40-250 µmol scale on NittoPhase UnyLinker® Solid Support (400 µmol/g) (40). Nucleoside phosphoramidites (fully protected) were incorporated under standard conditions: deblock with 3% dichloroacetic acid in dichloromethane; activate with 1 M 4,5-dicyanoimidazole and 0.1 M *N-*methylimidazole in acetonitrile; cap with 10% acetic anhydride in tetrahydrofuran (THF) and 10% *N*-methylimidazole in THF:pyridine; oxidize with 0.05 M iodine in 9:1 pyridine:water for phosphodiesters (PO), and 0.1 M xanthane hydride in 3:2 pyridine:acetonitrile (V:V) for phosphorothioates (PS). For all ASOs, and for siRNA sense strand, a 5’ hexylamino moiety was introduced with an MMT protected hexylamino phosphoramidite. All phosphoramidites were dissolved in 1:1 acetonitrile:toluene (v:v) at a concentration of 0.1 M. Coupling times for incorporation were 3 minutes for DNA phosphoramidites and 12 minutes for all other phosphoramidites. After synthesis, deprotection and cleavage from the resin were achieved by treating with 20% diethylamine in toluene for 20 minutes followed by suspension of the solid support in aqueous ammonia (28-30%). ASOs were incubated overnight at 55°C. siRNAs were incubated at room temperature for 48 hours. Filtration separated the solid support from the crude oligonucleotide, which was then purified with strong anion exchange chromatography (SAX) using Source 30Q resin (Cytiva, Cat. 17-1275-0). SAX buffers were A: 100 mM NH_4_OAc in 3:7 acetonitrile:water; B = 100 mM NH_4_OAc, 1.5 M NaBr in 3:7 acetonitrile:water. Purified oligonucleotides were desalted by HPLC on a C18 reverse-phase on a column (Waters, C-18 SPE, Cat. WAT043345) and lyophilized. Oligonucleotides synthesized for this work were mouse Malat1 3-10-3 2′-constrained ethyl (cEt) gapmer ASO full phosphorothioate backbone, sequence: 5’-GCATTCTAATAGCAGC-3’; mouse Malat1 5-10-5 2′-*O*-methoxyethyl (MOE) gapmer ASO mixed backbone (phosphodiester at linkages 2-5, 16-17, phosphorothioate at all other linkages), sequence: 5’-GCCAGGCTGGTTATGACTCA-3’; mouse and NHP HPRT1 siRNA antisense strand: MOE-5’-(*E*)-vinylphosphonate at position 1, 2’-F RNA at positions 2, 6, 14 and 16, 2’-OMe-RNA at all other positions, phosphorothioate at linkages 1-2, 21-22, phosphodiester at all other linkages, sequence: 5’-TUAAAAUCUACAGUCAUAGGAAU-3’; mouse and NHP HPRT1 siRNA sense strand: 2’-F-RNA at positions 7, 9-11, 2’-OMe-RNA at all other positions, phosphorothioate at linkages 1-2, 19-20, phosphodiester at all other linkages, sequence: 5’-UCCUAUGACUGUAGAUUUUAA-3’; NHP MAPT siRNA antisense strand: MOE-5’-(*E*)-vinylphosphonate at position 1, 2’F-RNA at positions 2, 14 and 16, MOE at positions 22-23, β-D-xylose at position 7, 2’-OMe-RNA at all other positions, phosphorothioate at linkages 1-2, 21-22, phosphodiester at all other linkages, sequence: 5’-TGCCUAAUGAGCCACACUUGGAA-3’; NHP MAPT siRNA sense strand: 2’-F-RNA at positions 9-11, 2’-*O*-MOE at positions 1-2, 14, 20-21, 2’-OMe-RNA at all other positions, phosphorothioate at linkages 1-2, 19-20, phosphodiester at all other linkages, sequence: 5’-CCAAGUGUGGCUCAUUAGGCA-3’. Hexylamine at the 5’ end of ASOs and siRNA was linked via phosphodiester unless stated otherwise. siRNA duplexes were formed either immediately prior to conjugation (BCN conjugates) or after purification of sense strand conjugates (maleimide conjugates).

### Synthesis of 5’-BCN-hexylamino oligonucleotides and conjugation to VHH-Fc-N3

The 5′-hexylamino oligonucleotide was dissolved in aqueous sodium tetraborate (borate) buffer (100 mM, pH 8.5) to a final concentration of 100 mg/mL. BCN-NHS (Millipore Sigma, Cat. 744867) dissolved in DMF at a volume equal to that of the oligonucleotide solution was added at 3 molar equivalents relative to the oligonucleotide. The reaction mixture was stirred at room temperature for 30 minutes, and reaction completion was monitored by HPLC-MS analysis. Following completion, the reaction mixture was diluted with water (5 mL) and filtered through a 0.45 µm PTFE syringe filter. The oligonucleotides were purified by SAX HPLC using Source 30Q resin (Cytiva, Cat. 17-1275-03). The SAX mobile phases were Buffer A: 100 mM ammonium acetate in 30:70 acetonitrile/water; Buffer B: 100 mM ammonium acetate and 1.5 M sodium bromide in 30:70 acetonitrile/water. Fractions containing full-length oligonucleotide, as determined by LC-MS analysis, were pooled and lyophilized. The resulting material was loaded onto a 5 g C18 SPE cartridge (Waters, Cat. WAT043345), washed sequentially with 1 M NaCl and water, and eluted with 50:50 acetonitrile/water. The eluate was lyophilized to yield the 5′-BCN-conjugated oligonucleotides. For siRNA conjugation, 5′-BCN-conjugated siRNA sense strands (BCN-ssHprt) were resuspended in water to 10 mM and annealed with complementary antisense strands (asHprt). The duplex siRNA was then combined with VHH-Fc-N3 in PBS containing 50 mM EDTA to final concentrations of 0.08 mM BCN-ssHprt, 0.10 mM asHprt, and 0.10 mM VHH-Fc-N_3_. For ASO conjugation, BCN-conjugated ASOs were resuspended in water to 10 mM and mixed with VHH-Fc-N_3_ to final concentrations of 0.8 mM ASO and 1.0 mM VHH-Fc-N_3_. Conjugation reactions were incubated overnight at 37°C and purified by anion-exchange chromatography (AEX) using an ÄKTA fast protein liquid chromatography (FPLC) system equipped with a HiTrap Capto Q ImpRes column (Cytiva, Cat. 17-5470-55). Briefly, the crude reaction mixture was filtered through a 0.2 μm PES syringe filter and loaded onto the column. The column was washed with 10 column volumes of AEX Buffer A (1× PBS), and conjugates were eluted using a 20-column-volume linear gradient from AEX Buffer A to AEX Buffer B (2 M NaCl in 1× PBS). All purified conjugates were characterized by liquid chromatography–mass spectrometry (LC-MS).

### Synthesis of HPRT1 siRNA sense strand containing 5’-maleimidepropionyl (C_3_-maleimide) linker

To a solution of 5′-hexylamino oligonucleotide (1 µmol) in 50 mM sodium phosphate buffer (pH 7.0, 100 µL/µmol) was added a solution of *N*-succinimidyl 3-maleimidopropionate (5 µmol) in DMSO. The reaction mixture was stirred at room temperature for 2 hours and then diluted with water (2 mL/µmol). The crude product was purified by SAX HPLC using a Source 30Q column (Cytiva, 30 µm, 2.54 × 8 cm). Mobile phase A consisted of 100 mM ammonium acetate in 30% aqueous acetonitrile, and mobile phase B consisted of 1.5 M NaBr in mobile phase A. Purification was performed using a linear gradient from 0–60% B over 60 minutes at a flow rate of 14 mL/minute. Fractions containing the full-length oligonucleotide, as confirmed by LC–MS analysis, were pooled and diluted with water to get 10% acetonitrile solution and desalted by reverse-phase HPLC to afford the corresponding 5′-maleimidopropionyl (C_3_-maleimide)-conjugated oligonucleotide in 80–85% isolated yield.

### Ring opening of HPRT1 siRNA sense strand containing 5’-C_3_-maleimide open-PO-C5h19VHH-Fc

To a solution of 5’- C_3_-maleimide-C5h19VHH-Fc conjugate (17.41 mgs, 302.34 µM) in 1X PBS (0.82 mL) solid sodium bicarbonate was added to obtain a saturated solution (120 mg). To this, a solution of EDTA was added (3.3 µL of 0.5 M solution to get 2 mM EDTA). The resulting solution was vortexed for 5 minutes. Precipitate of undissolved sodium bicarbonate was visible. The reaction mixture was kept at room temperature for 5 days to yield C_3_-maleimide open-PO-C5h19VHH-Fc. An equimolar amount of antisense strand was then added, and the resulting material was purified by size exclusion chromatography.

### Maleimide oligonucleotide conjugation to VHH-Fc-SH

Maleimide conjugated siRNA sense strands (mal-ssHprt or mal-ssMAPT) were resuspended in 50 mM sodium citrate, pH 6.0 to 10 mM and mixed together with VHH-Fc-SH in PBS plus EDTA 50 mM at final concentrations of 0.30:0.10 mM (mal-ss:VHH-Fc-SH) and incubated overnight at room temperature. Maleimide conjugated ASO (mal-Malat1) was mixed together with VHH-Fc-SH in PBS plus EDTA 50 mM at final concentrations of 0.30:0.10 mM (mal-ASO:VHH-Fc-SH) and incubated for 20-40 hours at room temperature. All conjugates were purified by AEX FPLC as above. Maleimide-sense strand conjugates were duplexed with their corresponding antisense strands at a 1:1 molar ratio to form conjugated siRNAs. All conjugates were verified by LC-MS and quantified by UV absorbance at 260 nm.

### Expression and purification of VHH-Fc

For microbial transglutaminase (MTGase) labeled VHH-Fc heterodimers, plasmids encoding monomeric VHH-Fc and monomeric Fc were co-transfected in ExpiCHO S cells (Thermo Fisher Scientific, Cat. A29133) according to the manufacturer’s recommendations. For thiol labeled heterodimers, plasmids encoding monomeric VHH-Fc and monomeric Fc were transfected and purified separately. All transfections were performed in the same fashion. Briefly, 50 mL CHO cultures were transfected with 40 µg total plasmid and cultured in a humidified incubator at 37°C, 8% CO_2_ on a shaking platform set to 140 RPM (day 0). The subsequent day (day 1), cultures were fed and transferred to a humidified incubator at 32°C, 5% CO2 with shaking as above. Cultures were fed again (day 5) and returned to the same incubator until harvest. To harvest antibody-containing supernatant, cells and debris were removed by centrifugation at 7,000 g for 10 minutes at 4°C (day 12). Supernatants were flocculated (41) with PEG6000 (3% w:v) (Rigaku Reagents, Cat. 101443-874) and Poly(diallyldimethylammonium chloride) (PDADMAC) MW 400,000-500,000 (Millipore Sigma, Cat. 409030) and incubated for 1 hour at room temperature followed by 1 hour on ice. To finalize clarification, a second centrifugation step at 7,000 g for 30 minutes at 4°C was followed by vacuum filtration through a 0.2 um PES filter. Antibodies were purified from crude supernatants by protein A affinity chromatography (MabSelect PrismA; Cytiva Cat. 17549852) on an ÄKTA Pure 25 FPLC (Cytiva, Cat. 29018224) at 4°C. Briefly, clarified supernatants were applied to the column, washed with 10 column volumes (CV) of PBS, washed with 10 CV of PBS plus 500 mM NaCl, and eluted with 10 CV of elution buffer (sodium chloride 100 mM, sodium acetate 100 mM, pH 3.5). Eluate fractions were neutralized with 1/5 fraction volume of Tris 1M, pH 8.0 and buffer exchanged to PBS or HEPES buffered saline (HBS) (HEPES pH 7.5, 25 mM; NaCl, 150 mM) by dialysis at 4°C overnight. Following buffer exchange, antibodies were quantified by UV absorbance at 280 nm using molar extinction coefficients (E280, M^-1^cm^-1^) and molecular weights estimated from primary amino acid sequences using ProtParam (42). Estimated protein extinction coefficients at 260 nm (E260) were derived by multiplying E280 by A260/A280 from the corresponding UV spectra. VHH-Fc scale-up for NHP studies was performed by Wuxi Biologics.

### Preparation of VHH-Fc-N_3_ by microbial transglutaminase

For azide addition via MTGase, antibodies were engineered with either a single monovalent C-terminal Myc (EQKLISEEDL) tag, internal MTGase (LLQG at positions 294-297) tag (bivalent) or C-terminal MTGase (LLQGPA) tags (monovalent or bivalent). Activa TI MTGase (Modernist Pantry, Cat. 1203) was reconstituted in PBS at 12.6 U/mL (approximately 0.2 U/mg dry powder). Antibody, 11-azido-3,6,9-trioxaundecan-1-amine linker (Millipore Sigma, Cat. 17758), and MTGase solutions were added together and diluted to final concentrations of 30 µM (antibody conjugation sites), 7.5 mM (250 linkers:1 conjugation site), and 7.04 U/mL, respectively. The reaction was stirred gently at room temperature for 1 hour. VHH-Fc-N3 was purified from MTGase and excess linker by Protein A chromatography, buffer exchanged to PBS or HBS and quantified exactly as described previously. Azide labeling for NHP studies was performed by Wuxi XDC.

### Preparation of VHH-Fc-SH for maleimide and bromoacetamide conjugation

Homodimeric VHH-Fc(Knob) bearing engineered cysteine (S239C or E388C) and homodimeric Fc-(Hole) were expressed, purified, and buffer exchanged to PBS as outlined previously. To uncap engineered cysteines and to generate heterodimers, equimolar amounts of VHH-Fc(Knob) and Fc(Hole) were combined together with 100 equivalents of reducing agent, either dithiothreitol (DTT) (Gold Biotechnology, Cat. DTT10) or tris-(2-carboxyethyl) phosphine (TCEP) (Gold Biotechnology, Cat. TCEP10) in PBS plus 5 mM EDTA (Thermo Fisher Scientific, Cat. 15575020) and stirred gently overnight at room temperature. Reactions were diluted 10-fold in cation exchange (CEX) Buffer A (sodium succinate, pH 5.0, 20 mM) and purified via FPLC on a HiTrap CaptoS ImpAct CEX column (Cytiva, Cat. 17371755). Excess reducing agent was removed by washing with 10 column volumes of CEX Buffer A and protein was eluted in 5 column volumes of CEX Buffer B (Tris, pH 7.5, 20 mM plus NaCl 150 mM). Hinge disulfides were then reoxidized by treatment with 50 equivalents dehydroascorbic acid (DHAA) (Millipore Sigma, Cat. 261556) for 3 hours at room temperature. For small volumes, excess DHAA was removed by 4 rounds of ultrafiltration and buffer exchange as described previously. For larger volumes, excess DHAA was removed by 3 rounds of dialysis as described previously. Following dialysis, VHH-Fc-SH heterodimers were quantified by A280 as described previously.

### Preparation of VHH-Fc-N_3_ from VHH-Fc-SH

For azide addition to VHH-Fc-SH, deblocked heterodimeric VHH-Fc-SH was incubated with 10 equivalents of bromoacetamido-PEG_2_-azide (Broadpharm, Cat. BP-22868) for 1 hour at room temperature. For small volumes, excess linker was removed by 4 rounds of ultrafiltration and buffer exchange into PBS through a 30 kDa molecular weight cutoff Amicon Ultra-4 (Millipore, Cat. UFC8030) or Ultra-15 (Millipore, Cat. UFC9030) centrifugal filter. For larger volumes, excess linker was removed by 3 rounds of dialysis (at least 8 hours per round) against at least 100 volumes of PBS in SnakeSkin dialysis tubing, 10 kDa MWCO (Thermo Fisher Scientific, Cat. 88245) with gentle stirring. Following linker removal, VHH-Fc-PEG_2_-N_3_ was quantified by A280 as described previously.

### Determination of the affinity of the VHH, VHH-Fc and VHH-Fc-siRNA conjugates to TfR1 proteins using bioluminescence resonance energy transfer (BRET)

Binding affinities of VHH, VHH-Fc, and VHH-Fc-siRNA conjugates to different species of TfR1 protein were evaluated using a nanoBRET (nBRET) assay. TfR1-nLuc fusion constructs were created by linking nanoLuc (nLuc) (Promega) through its N-terminal Val to the C-terminal F760 residue of human, cynomolgus, and mouse TfR1 via a GGGSGGSSG flexible linker and cloned into pLVX-IRES-Puro (Takara Clontech, Cat. 632183). HEK 293 cells stably expressing individual TfR1-nLuc proteins were then generated via lentiviral particle infection followed by puromycin selection. Crude membrane fractions from TfR1-nLuc expressing cells were resuspended in PBS and dispensed into white 96-well assay plates (Thermo Fisher Scientific, Cat. 136101) at 100μL per well at about 10,000 cell equivalents. To generate a tracer for competition assays, C5VHH with a C-terminal cysteine was labeled with Alexa Fluor 594 C_5_ maleimide reagent (Thermo Fisher Scientific, Cat. A10256), purified, and quantified according to manufacturer recommendations. To measure binding affinity of tracer for TfR1 proteins, 11.1 uL of C5VHH-594 was added to triplicate wells at 10x final concentration to produce a 12-point dilution series ranging from 1,000 to 0.006 nM, and incubated for 2.5 hours. To initiate BRET, 12.4 µL of 100 µM of the nLuc substrate furimazine (Promega, Cat. N157A) was added to each well and the mixtures were incubated for 5-30 minutes. The assay plate was then read on a Promega GlowMax Discover plate reader at wavelengths of 450 nm and 600 nm, and the ratio of emissions at wavelengths 450/600 was used to yield %BRET efficiency. Data was subjected to non-linear regression, then fitted to a single site binding hyperbolic function to derive the tracer Kd, the concentration of tracer at which 50% of maximum BRET ratio (Bmax) was achieved. All other binding affinities were determined by competition assays as follows: VHH and VHH-siRNA conjugates were titrated at 10x final concentration through a constant 10x concentration of C5VHH-594 tracer corresponding to 20x the Kd determined for the TfR1 species tested. Samples were incubated for 2.5 hours at room temperature to reach binding equilibrium followed by addition of furimazine and BRET measurement as described above. Inhibition constants (Ki) were obtained by fitting %BRET efficiency values to a competitive inhibition model, using the Kd value estimated for C5VHH-594 obtained in the same run. Values are presented as the average of triplicate data.

### VHH-Fc uptake in HEK293 and HEK293-TfR1 cells

HEK293 cells were cultured in DMEM (Thermo Fisher Scientific, Cat. 11965092) supplemented with 10% fetal bovine serum (Thermo Fisher Scientific, Cat. A5669801) in a humidified 37°C incubator with 5% CO_2_. To generate TfR1 overexpressing (HEK293-TfR1) cells, parental HEK293 were transduced with lentivirus bearing expression cassettes for human or mouse TfR1 and puromycin resistance. Cells were treated with puromycin dihydrochloride (Thermo Fisher Scientific, Cat. A1113803), 1 µg/mL for 72 hours or until non-transduced control cells were all dead. For cellular uptake studies, glass coverslips (Epredia, Cat. 22X22-1-001G) were immersed in 70% ethanol and allowed to dry fully. Coverslips were then coated with Poly-L-Lysine (PLL) (Millipore Sigma, Cat. P4832) for 30 minutes at room temperature. PLL was removed and coverslips were allowed to dry fully in 6-well cell culture dishes. Parental HEK293 and HEK293-TfR1 cells were grown to 70% confluency and dissociated with TrypLE (Thermo Fisher Scientific, Cat. 12605010). Cells were seeded on coverslips at 100,000 cells per well in 2 mL culture medium and returned to the incubator. Once cells reached 50-70% confluency (approximately 2 days post seeding), cells were fed C5VHH-Fc at 100 nM final concentration. Unfed wells served as negative controls. Following uptake, cells were washed 3 times with DPBS (Thermo Fisher Scientific, Cat. 14040133) and fixed with 4% formaldehyde (Thermo Fisher Scientific, Cat. 28906) in DPBS for 15 minutes at room temperature. Cells were then washed once with DPBS and permeabilized with 0.15% saponin (Millipore Sigma, Cat. 47036) in DPBS for 10 minutes at room temperature. Cells were again washed once with DPBS and blocked with 1% bovine serum albumin (BSA) (Millipore Sigma, Cat. A9418) in DPBS for 30 minutes at room temperature. Cells were then stained with FITC conjugated anti-human secondary antibody (Abcam, Cat. ab97224) diluted 1:100 in 1% BSA-DPBS for 1 hour at room temperature, washed 4 times with DPBS, washed once with water, and mounted with ProLong Diamond Antifade Mountant with DAPI (Thermo Fisher Scientific, Cat. P36971) on frosted microscope slides (Corning, Cat. CLS294875X25). Mounted slides were cured in the dark overnight at room temperature and were then imaged on an Olympus FV-1000 confocal microscope. Composite false color images were generated with ImageJ software.

### Generation of human TfR1 knock-in mice and mouse dosing

The human TfR1 knock-in (KI) mouse was generated as described in Østergaard et al (5). Briefly, CRISPR–Cas9–mediated genome editing was used to replace, within the mouse Tfrc locus, the coding sequence of exon 2 together with the intron 2 splice-donor site with the human TFRC open reading frame. Therefore, the resulting KI allele expresses human TFRC under the control of the endogenous mouse promoter. Homozygote human TfR1 KI mice, which express only the human and not the murine TfR1 were used. Experiments in adult wild-type C57BL/6NTac mice and in human TfR1 KI mice were conducted at Ionis Pharmaceuticals. Animals were assigned to groups (n = 3–4 per group) and acclimated for ≥7 days before dosing. Mice were housed in a virus-free barrier facility on a 12-hour light/dark cycle with ad libitum access to food and water. Dosing was performed by IV or SC injection, as specified in the main text and figure legends. Animals received phosphate-buffered saline (PBS; vehicle control), unconjugated siRNA/ASO (Supplementary Figure 3), or VHH-Fc-conjugated siRNA/ASO, as specified in the figures. The dose levels, which are reported in the figures, are expressed in mg/kg body weight and refer to the oligonucleotide portion of the moiety (ASO or siRNA equivalents). Unless otherwise indicated, mice were dosed on days 1 and 8, and were euthanized two weeks after the last dose, on day 22. In the experiment with C5VHH-Fc-Malat1 ASO (Figure 2D, E; Supplementary Figure 13B) the vehicle was HEPES instead of PBS, and the mice were euthanized one week after the last dose, on day 16. In the experiment described in Supplementary Figure 5, mice were dosed either by IV injection on days 1 and 8, and euthanized on day 22, or by SC injection on day 1 and euthanized on day 22. In the case of Supplementary Figure 15, the mice were pre-dosed with anti-mouse CD4 monoclonal antibody, a commonly used immunomodulatory strategy to attenuate adaptive immune responses—particularly anti-drug antibody (ADA) formation—in mice against biologics containing a human IgG Fc region (43–45). Specifically, 0.5 mg of the anti-mouse CD4 antibody (Clone YTS 177; BioXcell, Cat. BE0003-3) were administered via intraperitoneal injection on days 0 and 7.

All procedures in C57BL/6NTac and human TfR1 KI mice complied with the eighth edition of the Guide for the Care and Use of Laboratory Animals (AAALAC-accredited Unit 000962 and State of California Department of Public Health Certificate No.071) and were conducted under the Institutional Animal Care and Use Committee (IACUC) protocol 2023-1219.

### Evaluation of VHH-Fc siRNA conjugates in non-human primates

Cynomolgus macaques (*Macaca fascicularis*) of 2–6 years of age, were randomized to treatment groups (n = 3–4 per group). Animals received VHH-Fc-siRNA conjugates by either a 60-minute IV infusion (4 mL/kg volume) or subcutaneous bolus injection (0.2 mL/kg volume). The animals were dosed on study days 1, 8, 15, and 22 (four total doses), and were euthanized on either day 24 or day 36. For the study reported in Figure 4 and Supplementary Figure 11, one animal in each of the two IV treatment groups was dosed as a sentinel, approximately 48 hours before the remaining animals in those groups and was euthanized on day 38. At necropsy, tissues were collected, weighed, and either flash-frozen for RNA analyses or fixed in neutral-buffered formalin for histological evaluation. When knockdown data are reported for combined NHP cortical brain regions, the analysis includes frontal, motor, temporal, and occipital cortices. The NHP studies were conducted at Keyprime Research Co., Ltd. (KPR; Republic of Korea) and approved by the IACUC at KPR.

### Animal tissue homogenization and RNA extraction

Mouse tissues were excised and immediately placed into wells of 96-well plates (Corning, Cat. 29445-166) that had been prechilled on dry ice and maintained on dry ice throughout tissue collection, then stored at −80°C. For NHP tissue analysis, brains were sectioned into 4-mm coronal slices, quadriceps were sampled as 5–10-mm transverse mid-muscle slices, and hearts were cut longitudinally to open all 4 chambers; tissues were collected flat on a metal plate placed on dry ice, flash-frozen in liquid nitrogen, and stored at -80°C. At the time of sampling, biopsy punches were obtained from each frozen tissue slice; punches were collected from distinct anatomical regions of the brain, defined by morphology and spatial orientation. Different sections of the spinal cord, liver, and other NHP tissues were also dissected and flash-frozen in liquid nitrogen. Tissues were homogenized in TRIzol reagent (Thermo Fisher Scientific, Cat. 15596026). Total RNA was isolated by phenol–chloroform extraction and further purified using the RNeasy 96 Kit (Qiagen, Cat. 74181).

### Analysis of gene expression by quantitative real-time RT-PCR

Target transcript abundance was quantified in each sample by quantitative real-time RT-PCR (qRT-PCR) using the Express One-Step qRT-PCR kit (Thermo Fisher Scientific, Cat. 11781200) and a standard 2 hour cycling program, per the manufacturer’s manual. Forward and reverse primers, plus a fluorescently labeled probe with a quencher system were used. The primers-probe sets were obtained from Integrated DNA Technologies (IDT) except for the TaqMan™ NHP HPRT1 gene expression assay (Thermo Fisher Scientific, Cat. Rh02800695_m1). The sequence of primers and probes are listed in Supplementary Table 1. Gene-expression values were normalized to the housekeeping gene GAPDH and expressed as percent change relative to the appropriate control group(s).

### Statistical analysis

Statistical analyses were conducted using GraphPad Prism software. For comparisons involving more than one treatment group, statistical significance relative to vehicle control was assessed by one-way analysis of variance (ANOVA) followed by Dunnett’s multiple-comparisons post hoc test. When only two groups were compared, consisting of one treatment group and one vehicle control group, an unpaired Welch’s t-test was used. P values ≤ 0.05 were considered statistically significant and are denoted by an asterisk (*) in the figures. Except for Supplementary Figure 5, significance annotations indicate only differences between the vehicle control group and each treatment group; pairwise comparisons between different treatment groups were not performed and are therefore not reported. Error bars represent the standard deviation (SD).

## RESULTS

### TfR1-binding VHH molecules do not compete with transferrin and are taken up into TfR1 expressing cells

Recently, David et al. described two llama-derived VHH ligand families, C5 and B8, that bind to human, mouse, and NHP TfR1 and cross the BBB in mice (46). With an eye toward clinical translation, we sought to explore the utility of these VHH families for the delivery of oligonucleotides to the CNS of rodents and NHPs.

Our initial studies examined binding of VHH family members to human, mouse, and NHP TfR1, as well as lack of competition with Tf. To that end, we expressed C5VHH in CHO cells and measured its binding affinity to human, mouse, and NHP TfR1 via nBRET assay, observing binding to all three proteins (Figure 1A). The equilibrium binding affinities (Kd) were 0.27 nM for human TfR1, 16.53 nM for NHP TfR1, and 11.30 nM for mouse TfR1 (Figure 1A).

**Figure 1:**
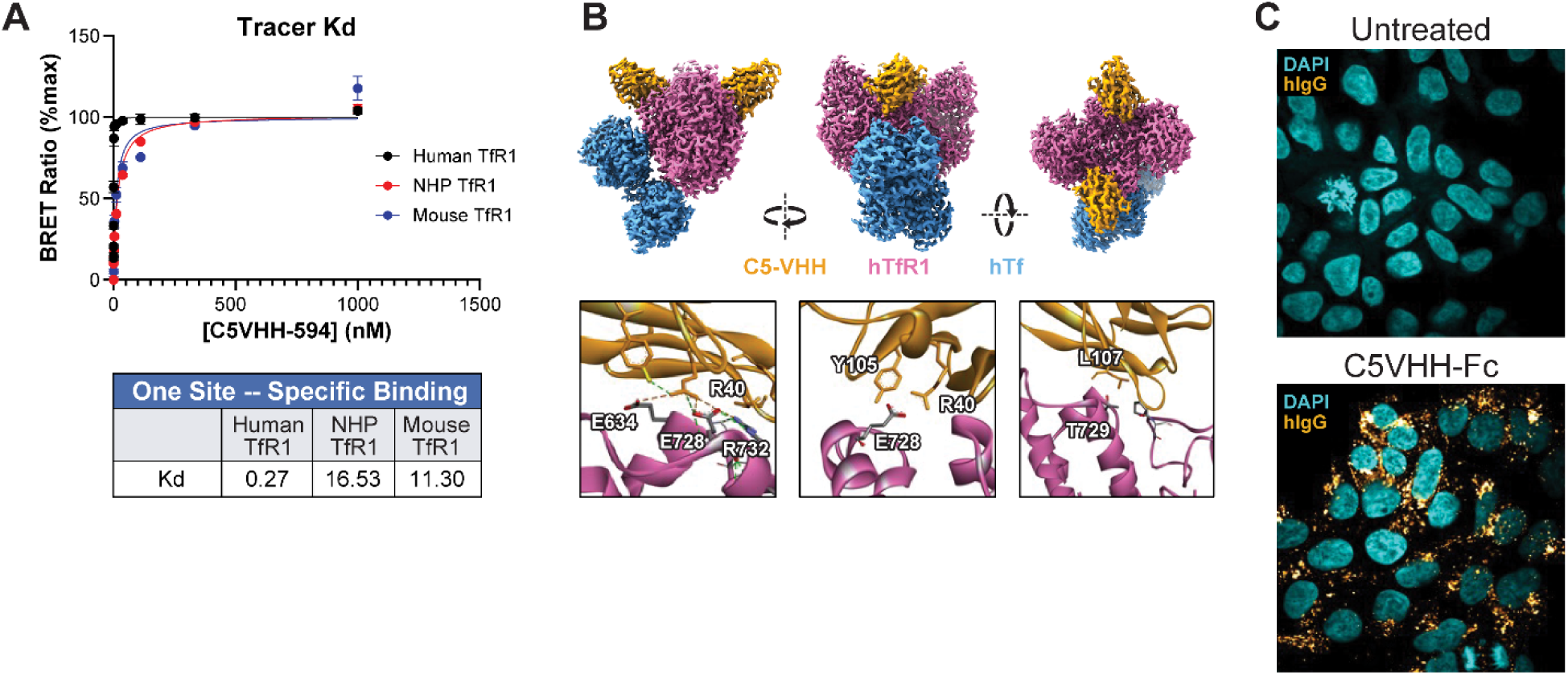
TfR1-binding VHH molecules do not compete with transferrin and are taken up into TfR1 expressing cells. (**A**) nBRET binding of C5VHH to human, NHP (*Macaca fascicularis*), and mouse TfR1. (**B**) Cryo-EM structure of C5VHH (gold) in complex with human TfR1 (hTfR1, magenta) and human holo-Tf (hTf, blue). Inset panels detail sidechain interactions between C5VHH (R40, Y42, Y105, L107) and human TfR1 (P323, E634, E728, T729, R732). (**C**) Confocal micrographs of HEK293 cells expressing mouse TfR1, either untreated or treated with C5VHH-Fc, and stained with DAPI (cyan) and anti-hIgG-FITC (gold).

Previous reports suggest that C5VHH does not compete with Tf for binding to TfR1 (46). To fully elucidate the binding site of C5VHH on TfR1, we performed cryo-EM on the complex of human TfR1 with holo-Tf and C5VHH. We observed C5VHH binding to a conformational epitope spanning the two TfR1 monomers adjacent to, but not overlapping with, the Tf binding site (Figure 1B). Upon examination of the structure, we observed interactions of C5VHH residues R40, Y42, Y105 and L107 (IMGT numbering) with TfR1 residues E634, G636, E728, and R732 (Figure 1B), supporting the reported role (46) of these TfR1 residues in mediating C5VHH binding.

To improve exposure to endothelial cells at the blood-brain interface, we sought to prolong the circulation half-life of VHH ligands by fusion to IgG1 Fc domains, which recycle IgG molecules following fluid-phase endocytosis. We employed human IgG1 Fc engineered with reduced immune effector functions (“LALA”: L234A, L235A (47) or “LALAPG”: L234A, L235A, P329G (48)) and “knobs-into-holes” heterodimerization (49, 50), where only one chain incorporates an N-terminal anti-TfR VHH. Finally, we appended a C-terminal tag for conjugation with MTGase to enable facile site-selective incorporation of bio- orthogonal conjugation handles (51–54). The monovalent anti-TfR1 VHH-Fc (VHH-Fc) was taken up by HEK293 cells overexpressing mouse or human TfR1. Following a short incubation, C5VHH-Fc puncta were observed in TfR1 overexpressing cells, but not in parental HEK293, which express a lower level of TfR1. (Figure 1C; Supplementary Figure 1A).

### Conjugation to VHH-Fc fusion proteins enables siRNA and ASO activity in CNS of wild-type and human TfR1 KI mice after systemic administration

We next labeled monovalent VHH-Fc proteins with a bifunctional amino-PEG_3_-azide linker via MTGase (Supplementary Figure 1B) and conjugated a BCN-siRNA that targets human, mouse, and NHP HPRT mRNA (Supplementary Figure 1C). Binding affinities of unconjugated VHH-Fc and the corresponding siRNA conjugates for TfR1 were measured by nBRET (Supplementary Table 2). The conjugates showed

binding affinities that were generally similar to, even if in some cases weaker or stronger than, those of unconjugated VHH-Fc (Supplementary Table 2). Specifically, C5VHH-Fc has binding affinities of 0.8 nM, 11.9 nM, and 11.3 nM for human, NHP, and mouse TfR1, respectively (Supplementary Table 2; Figure 1A), whereas the C5VHH-Fc-Hprt siRNA conjugate has binding affinities of 0.4 nM, 9.9 nM, and 24.5 nM for human, NHP, and mouse TfR1, respectively (Supplementary Table 2).

To determine whether VHH-Fc conjugation enables BBB crossing of antisense molecules in vivo, we dosed wild-type mice intravenously with the C5VHH-Fc-Hprt siRNA conjugate at 1 mg/kg (siRNA equivalents), and we measured its activity in CNS tissues. This dose was selected based on prior evidence that TfR1-targeted transport vehicles can enable systemic CNS delivery of therapeutic oligonucleotides across the BBB (55). Notably, the affinity of C5VHH-Fc-Hprt siRNA for mouse TfR1, as measured by the nBRET assay, is 24.5 nM, whereas its affinity for human TfR1 is considerably higher (0.4 nM) (Supplementary Table 2). qRT-PCR analysis revealed that IV injection of C5VHH-Fc-Hprt siRNA at 1 mg/kg (siRNA equivalents) results in robust Hprt mRNA knockdown in the brain cortex and spinal cord of wild-type mice (Figure 2A), without evidence of BBB disruption, as demonstrated by the absence of fibrinogen staining in brain histological sections (Supplementary Figure 2). In contrast, the unconjugated Hprt siRNA dosed in mice intravenously is inactive in the CNS, even at the relatively high dose level of 10 mg/kg (Supplementary Figure 3A). Subcutaneous dosing of C5VHH-Fc-Hprt siRNA resulted in target mRNA reduction in murine CNS tissues comparable to that achieved by IV administration (Supplementary Figure 4). By virtue of TfR1 binding, C5VHH-Fc-Hprt siRNA knocked down Hprt mRNA very efficiently in skeletal (quadriceps) and cardiac muscles of wild-type mice (Figure 2A). Conversely, the unconjugated Hprt siRNA had poor potency in quadriceps, and especially in the heart (Supplementary Figure 3A). Interestingly, when dosed in human TfR1 KI mice by IV injection, the C5VHH-Fc-Hprt siRNA resulted in significant, yet notably inferior CNS activity compared to wild-type mice (Figure 2B). The lower target knockdown efficiency measured in brain cortex and spinal cord of the human TfR KI mice was associated with a much higher binding affinity of the C5VHH-Fc conjugate for the human TfR1 (0.4 nM) compared to the mouse receptor (24.5 nM), as measured by the nBRET assay (Supplementary Table 2). On the other hand, the Hprt mRNA reduction in skeletal and cardiac muscles was comparable between human TfR1 KI and wild-type mice (Figure 2A, B). In agreement with the qRT-PCR results, immunohistochemistry to detect HPRT protein in histological sections of the brain from the wild-type mice revealed robust, widespread target protein reduction across the mouse brain by the IV dosed C5VHH-Fc-Hprt siRNA (Figure 2C), but not the unconjugated siRNA (Supplementary Figure 3B).

**Figure 2:**
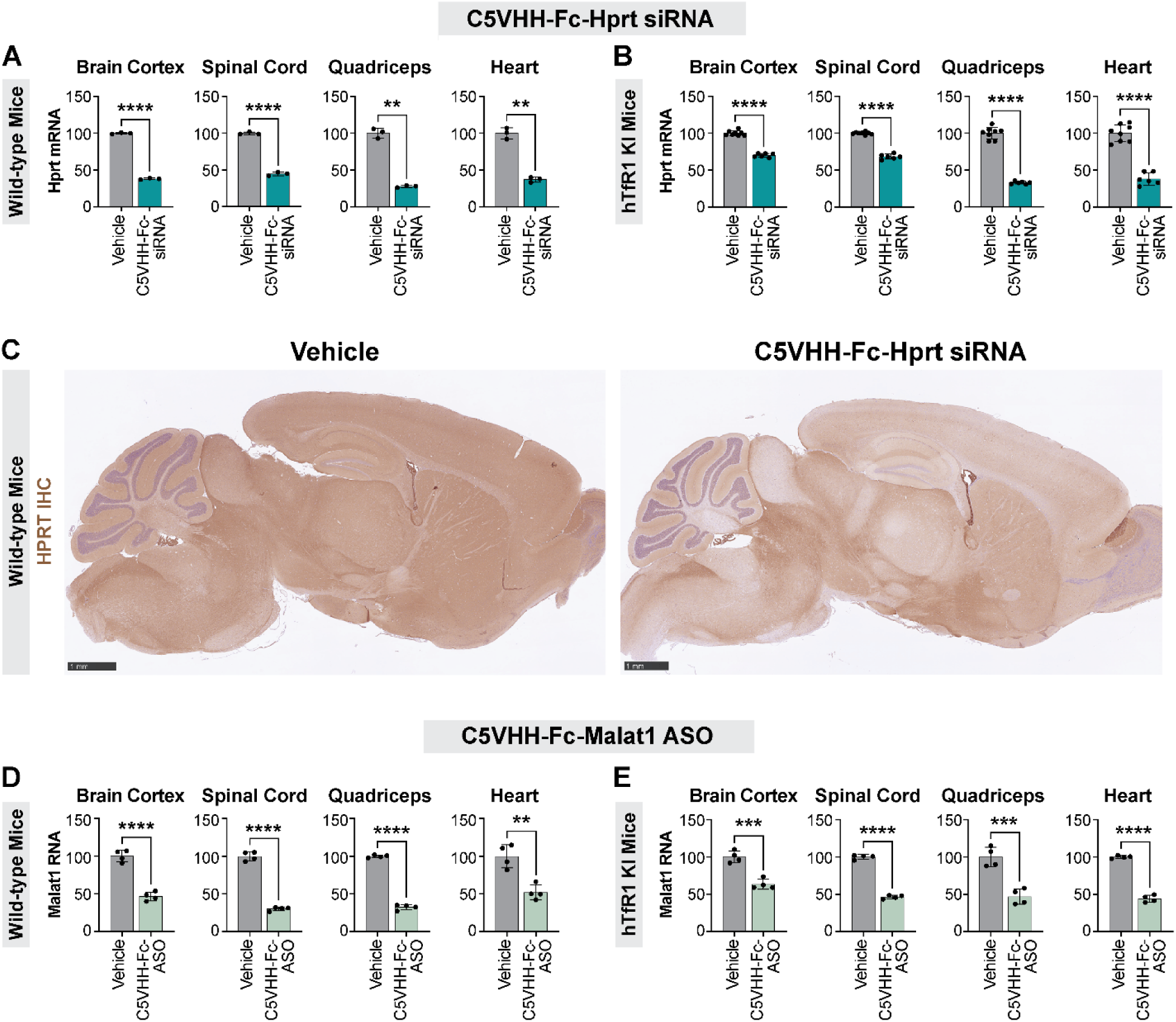
Conjugation to C5VHH-Fc enables siRNA and ASO activity in CNS of wild-type and human TfR1 KI mice following IV dosing. (**A**, **B**) qRT-PCR analysis for Hprt mRNA in tissues from (**A**) wild-type mice or (**B**) human TfR1 KI (hTfR1 KI) mice dosed IV with either vehicle or 1 mg/kg (siRNA equivalents) of C5VHH-Fc-Hprt siRNA on days 1, 8, and euthanized on day 22. (**C**) Immunohistochemistry (IHC) to detect HPRT protein (brown stain) in histological sections of the brain of wild-type mice (same mice as in A). Scale bars: 1 mm. (**D**, **E**) qRT-PCR analysis for Malat1 RNA in tissues from (**D**) wild-type mice or (**E**) human TfR1 KI mice dosed IV with either vehicle or 1 mg/kg (ASO equivalents) of C5VHH-Fc-Malat1 ASO on days 1, 8, and euthanized on day 16.

Similarly to siRNA, conjugation of a 16-mer full PS cEt-modified gapmer Malat1 ASO to C5VHH-Fc (Supplementary Figure 1D) resulted in robust target RNA knockdown in brain cortex, spinal cord, skeletal muscle and heart after IV dosing at 1 mg/kg (based on ASO equivalents) in wild-type (Figure 2D) and human TfR1 KI mice (Figure 2E). On the other hand, the unconjugated Malat1 ASO dosed IV in mice is inactive in brain cortex and is only modestly active in quadriceps muscle and heart (Supplementary Figure 3C).

We next asked whether increasing the ratio of siRNA cargo to antibody from a drug-antibody ratio (DAR) of 1 (DAR1) to 2 (DAR2) would improve CNS activity of VHH-Fc-siRNA conjugates. To that end, we dosed wild-type mice with DAR1 or DAR2 molecules, comprising one or two siRNAs conjugated to C5VHH-Fc, respectively (Supplementary Figure 1C). Mice received either two intravenous doses of 1, 3.5, or 10 mg/kg siRNA equivalents, or a single subcutaneous dose of 2, 7, or 20 mg/kg siRNA equivalents. These regimens were selected to keep the total amount of administered compound constant across routes over the course of the experiment. IV and SC administration were previously shown to produce similar knockdown efficiency (Supplementary Figure 4). qRT-PCR analysis to measure target Hprt mRNA levels in brain cortex and spinal cord revealed reduced activity, on an siRNA mg/kg basis, for DAR2 conjugates compared to DAR1, independently of the route of administration (Supplementary Figure 5). IV dosed DAR1 and DAR2 molecules perform similarly in the skeletal and cardiac muscles, whereas DAR2 loses some activity compared to DAR1 in the heart after SC injection (Supplementary Figure 5). In the liver, DAR2 is more potent than DAR1 (Supplementary Figure 5). For DAR1 and DAR2 conjugates, a single SC bolus administration was less effective than two IV injections in all tissues except quadriceps, where saturation prevented this comparison (Supplementary Figure 5).

### The CNS activity of IV dosed VHH-Fc-siRNA conjugates correlates with conjugate TfR1 binding affinity in wild-type and human TfR1 mice

Following IV delivery of C5VHH-Fc-conjugated Hprt siRNA (Figure 2A, B) and Malat1 ASO (Figure 2D, E) we observed lower CNS activity in human TfR1 KI mice (Figure 2B, E) compared to wild-type mice (Figure 2A, D).

Our interest lies in the potential therapeutic application of VHH-Fc-conjugated antisense molecules. Therefore, we sought to improve the human TfR1 mediated transcytosis of VHH-Fc-siRNA conjugates via selection of VHH ligands with optimal binding affinities for the human TfR1. Accordingly, we intiated a campaign of affinity maturation and humanization, generating five VHH variants of the C5 clone family with a variety of binding affinities for human TfR1 (Supplementary Table 2). To further understand the relationships between the TfR1 binding affinity and the BBB crossing properties of VHH-Fc-based siRNA conjugates, we undertook a systematic approach where the five different VHH clones (C5, C5h9, C5h19, C5h20, C5V30) were formatted as monovalent VHH-Fc (Supplementary Figure 1E), conjugated to Hprt siRNA, and dosed by IV injection at 1 mg/kg (siRNA equivalents) in wild-type (Figure 3A; Supplementary Figure 6A) and, most importantly, human TfR1 KI mice (Figure 3B; Supplementary Figure 6B). The activity of each construct was measured by qRT-PCR in brain cortex and expressed as percent reduction of the target Hprt mRNA compared to vehicle-treated mice (Supplementary Figure 6A, B). The target knockdown was then plotted against the binding affinity (nBRET Ki) towards the cognate TfR1 receptor species for each VHH-Fc-siRNA conjugate (Figure 3A, B; Supplementary Table 2). VHH-Fc-siRNA variants with TfR1 binding affinities ranging from high single-digit nM to low hundreds nM exhibited the strongest CNS activity, achieving more than 60% Hprt mRNA reduction in brain cortex after IV dosing (Figure 3A, B; Supplementary Table 2). Conversely, when the TfR1 binding affinity was either too high (low single-digit nM or sub-nM) or too low (several hundreds nM), the CNS activity of the IV-dosed VHH-Fc-siRNA variants was markedly reduced (Figure 3A, B; Supplementary Table 2).

**Figure 3:**
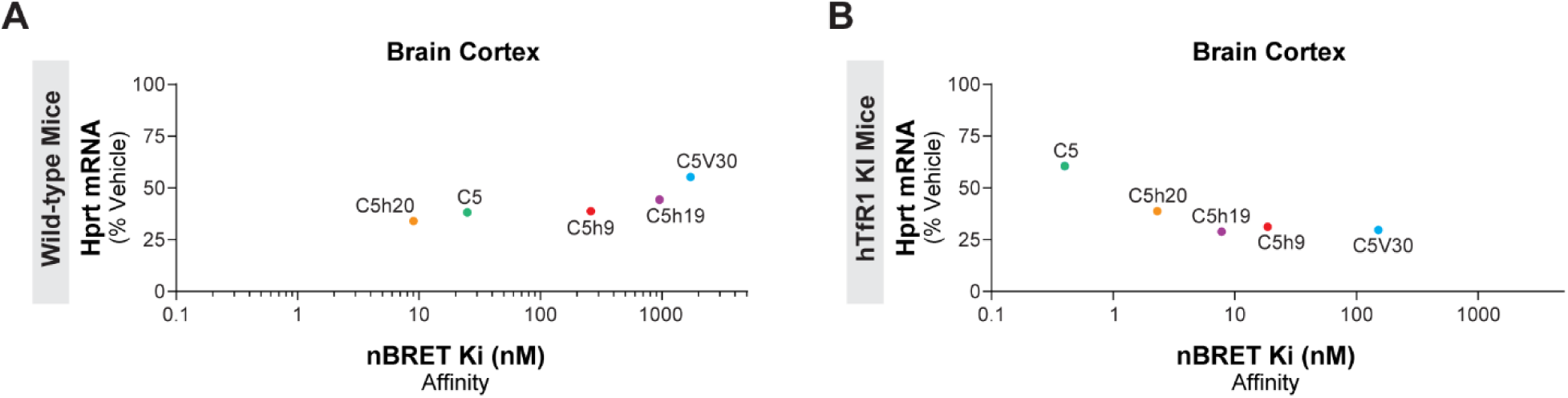
The binding affinity of the VHH ligand for the cognate TfR1 correlates with the CNS activity of VHH-Fc-siRNA conjugates after IV dosing in mice. (**A**) The target Hprt mRNA knockdown in brain cortex of wild-type mice (which express only mouse TfR1) was measured by qRT-PCR after IV dosing of various VHH-Fc-Hprt siRNA conjugates at 1 mg/kg (siRNA equivalents). The Hprt mRNA level is expressed as percentage of vehicle control animals and plotted versus the binding affinity (Ki) of each VHH-Fc-siRNA conjugate for the mouse TfR1, as measured by nBRET assay. The VHH clone is indicated, since it constitutes the part of the molecule that binds TfR1. (**B**) Similarly to A, the Hprt mRNA knockdown after IV dosing at 1 mg/kg in human TfR1 KI homozygote mice (which express only human TfR1) is plotted versus the binding affinity of the C5-series VHH conjugates for the human TfR1.

We observed that the range of TfR1 binding affinities required for maximum CNS activity appeared relatively broad in human TfR1 KI mice, extending in the hundreds nM (Figure 3B). This prompted us to dose the conjugates at the lower dose level of 0.3 mg/kg (siRNA equivalents), in an attempt to better differentiate the various VHH clones. Ultimately, the lower dose level experiment in human TfR1 KI mice confirmed that VHH-Fc-siRNA variants with TfR1 binding affinities ranging from high single-digit nM to low hundreds nM result in similar, very robust target knockdown in brain cortex after IV dosing (Supplementary Figure 7A, B). The activity of the different VHH-Fc-siRNA variants in the spinal cord followed a similar ranking as the brain cortex (Supplementary Figure 6A, B; Supplementary Figure 7B). On the other hand, the Hprt mRNA knockdown achieved in quadriceps muscle, heart, and liver was comparable across the five different VHH conjugates (Supplementary Figure 6A, B; Supplementary Figure 7B).

In addition to the C5-series, we also explored the relationship between CNS activity and TfR1 binding affinity for a closely related family of VHHs, called B8-series. Our goal is to identify optimal human TfR1 VHH ligands, so we screened fifteen different B8-series VHH-Fc-Hprt siRNA conjugates (Supplementary Figure 1E) with a broad range of binding affinities for human TfR1 (Supplementary Table 2), in human TfR1 KI mice. Nevertheless, we also dosed the fifteen compounds in wild-type mice, allowing us to expand our activity-affinity dataset by covering the Ki values of the same B8VHH ligands towards the mouse TfR1 (Supplementary Table 2). Several of the fifteen B8-series VHH-Fc-Hprt siRNA conjugates achieved robust Hprt mRNA knock down in brain cortex and in spinal cord of wild-type (Supplementary Figure 8A) and especially human TfR1 KI mice, after IV dosing at 1 mg/kg (siRNA equivalents) (Supplementary Figure 8B). Consistent with the C5-series data, the screening of B8-series conjugates in wild-type mice confirmed that binding affinities for TfR1 in the low tens nM are associated with strong CNS activity (Supplementary Figure 9A). In human TfR1 KI mice, several B8-series VHH conjugates led to similarly robust target knockdown in the CNS despite their wide range of binding affinities for human TfR1 (Supplementary Figure 9B). In an attempt to further differentiate the B8VHH clones, we injected additional human TfR1 KI mice with the B8-series VHH-Fc-Hprt siRNA conjugates at the lower dose level of 0.3mg/kg (siRNA equivalents). Such lower dose strategy marginally improved the ranking of the conjugates based on CNS activity, with lower binding affinities (mid-high hundreds nM) towards human TfR1 resulting in a sharper loss of CNS activity (Supplementary Figure 10A, B). Similar to what was observed with the C5-series, the activity of the B8-series VHH conjugates in skeletal muscle (quadriceps) and heart of both wild-type and human TfR1 KI mice was comparable across the fifteen compounds, despite their different binding affinities for TfR1 (Supplementary Figure 8A, B; Supplementary Figure 10B). In the liver, especially in human TfR1 KI mice, lower binding affinities were associated with less activity (Supplementary Figure 8A, B; Supplementary Figure 10B).

Taken together, our results suggest that TfR1 binding is a key determinant of BBB crossing and VHH-Fc-siRNA conjugate activity in the CNS following systemic delivery in vivo.

### Systemic dosing of C5VHH-Fc-Hprt siRNA results in robust target mRNA knockdown in the NHP CNS

Preclinical testing in NHP offers valuable translational validation of potential therapeutic molecules before advancing them to clinical trials. Accordingly, our ligand optimization was tailored towards the identification of NHP-human TfR1 cross-reactive VHHs, and utilized both the C5 and B8 families of VHH clones (Supplementary Table 2). In particular, the binding affinities of B8V40, B8V32h14, and B8V31h4 for the NHP TfR1 are within approximately 2-fold of the affinity for the human TfR1 (Supplementary Table 2). When dosed IV at 1 mg/kg (siRNA equivalents) in NHPs, HPRT siRNA conjugated to B8V40-Fc, B8V32h14-Fc, and B8V31h4-Fc (Supplementary Figure 1F) achieved significant target HPRT mRNA reduction in brain cortical regions (Supplementary Figure 8C; Supplementary Figure 9C), and spinal cord (Supplementary Figure 8C) compared to vehicle-dosed animals. Interestingly, compared to human TfR1 KI mice, in NHP we observed a much more rapid reduction in activity as the binding affinity decreased (Supplementary Figure 9B, C). On the other hand, all three B8V40-Fc-, B8V32h14-Fc-, and B8V31h4-Fc-HPRT siRNA conjugates were able to knock down HPRT mRNA more than 60% in NHP skeletal muscle (quadriceps) and heart, while mostly sparing the liver (Supplementary Figure 8C).

The C5VHH-Fc-HPRT siRNA conjugate has a binding affinity for NHP TfR1 of 9.9 nM (as measured by nBRET assay) (Supplementary Table 2). According to our studies in wild-type and human TfR1 KI mice, this type of affinity is within the optimal range for efficient RMT across the BBB. Indeed, when dosed in NHPs intravenously at 1 and 5 mg/kg (siRNA equivalents), C5VHH-Fc-HPRT siRNA (Supplementary Figure 1C) achieved excellent target mRNA knockdown across various brain regions, particularly in deeper structures such as the caudate and putamen (Figure 4A; Supplementary Figure 11). A notable exception is the cerebellar cortex, where the target knockdown was markedly lower relative to other brain regions (Supplementary Figure 11). Consistent with the qRT-PCR results, immunohistochemical analysis of NHP brain sections revealed uniformly reduced HPRT protein staining after IV administration of 5 mg/kg (siRNA equivalents) C5VHH-Fc-HPRT siRNA (Figure 4B). These findings were further supported by Western blot analysis of HPRT protein in the brain cortex and caudate, where robust target protein reduction was measured following IV dosing with 5 mg/kg C5VHH-Fc-HPRT siRNA (Supplementary Figure 12A, B).

**Figure 4:**
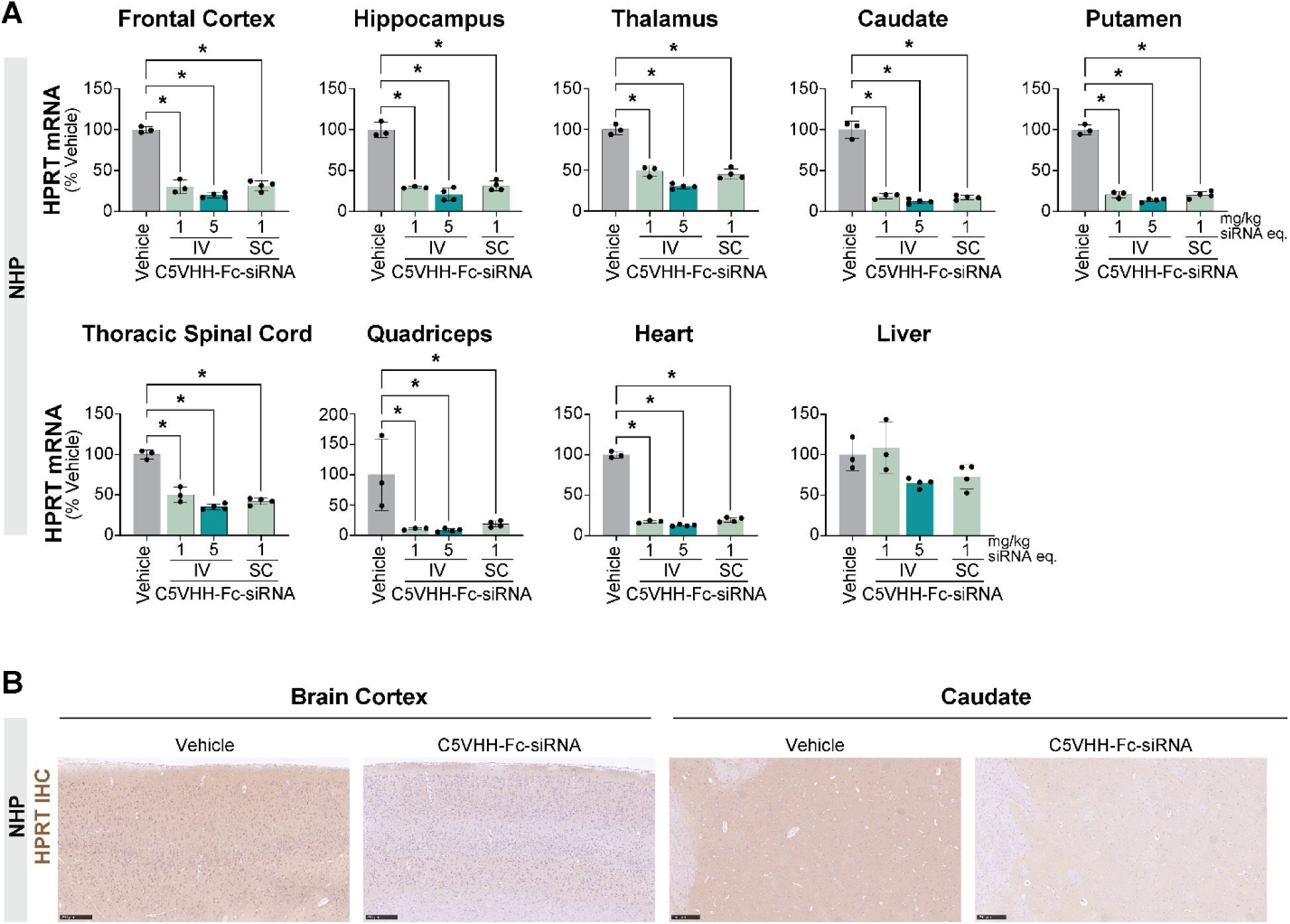
IV and SC dosing of C5VHH-Fc-HPRT siRNA results in robust target mRNA knockdown in the NHP CNS. (**A**) Target HPRT mRNA knockdown measured by qRT-PCR in various tissues from NHPs dosed IV or SC with C5VHH-Fc-HPRT siRNA at the indicated dose levels, based on siRNA equivalents (eq.). N equals 3 for the vehicle group and the 1 mg/kg IV group. N equals 4 for the 5 mg/kg IV group and the 1 mg/kg SC group. IV: dosing by intravenous infusion; SC: dosing by subcutaneous injection. (**B**) Immunohistochemistry (IHC) to detect HPRT protein (brown stain) in histological sections of brain cortex and caudate from NHPs dosed IV with either vehicle or C5VHH-Fc-HPRT siRNA at 5 mg/kg siRNA equivalents. Scale bars: 250 µm.

The HPRT mRNA knockdown measured by qRT-PCR in quadriceps muscle and heart from the NHPs dosed IV with 5 mg/kg C5VHH-Fc-HPRT siRNA reached 92% and 87%, respectively, compared to vehicle-dosed animals (Figure 4A). On the other hand, the liver was partially spared, with a modest 35% target HPRT mRNA knockdown measured (Figure 4A). Notably, C5VHH-Fc-HPRT siRNA dosed IV appeared to approach almost-complete knockdown, with target mRNA reduction already nearing a plateau at 1 mg/kg, such that the 5 mg/kg dose provided only marginally greater activity (Figure 4A).

Remarkably, subcutaneous injection of 1 mg/kg C5VHH-Fc-HPRT siRNA in NHP also achieved excellent target HPRT mRNA knockdown, comparable to the IV route of administration (Figure 4A; Supplementary Figure 11).

Interestingly, the C5VHH-Fc-HPRT siRNA conjugate was able to knock down its target mRNA also in dorsal root ganglia (DRG) and peripheral nerve in NHP (Supplementary Figure 13A). These observations, taken together with the robust, dose-responsive target Malat1 RNA reduction measured by qRT-PCR in DRG and sciatic nerve from wild-type mice dosed IV with C5VHH-Malat1 ASO (Supplementary Figure 13B), suggest that the VHH-Fc system can mediate RMT not only across the BBB but also the blood-nerve barrier (BNB).

### Optimization of conjugation site, but not linker chemistry, improves the CNS activity of systemically dosed VHH-Fc siRNA and ASO conjugates

Generation of antibody-drug conjugates (ADCs) has traditionally relied on chemical conjugation to naturally occurring, solvent accessible lysine or cysteine residues using simple and robust chemistries, including NHS-ester or maleimide (56). Because these reactions occur at multiple accessible residues, the resulting conjugates are inherently stochastic, yielding heterogeneous populations that differ in both DAR and conjugation site (57). Accumulating evidence from preclinical studies indicates that site-specific conjugation, beyond enabling precise control over DAR and attachment site, can favorably influence ADC pharmacokinetics, potency, and tolerability (58, 59). In particular, the choice of conjugation site itself has been shown to exert a substantial impact on these parameters (52, 54, 59). One widely used strategy to achieve site-specific conjugation is the introduction of engineered cysteine residues through protein engineering (58, 60, 61). Building on this premise, we explored two alternative conjugation sites for linking ASOs or siRNAs to VHH–Fc fusion proteins. To enable cysteine–maleimide (Cys-Mal) chemistry, we redesigned the VHH–Fc constructs by removing the C-terminal transglutaminase LLQGPA tag and introducing engineered cysteine residues at positions 239 or 388 (IMGT numbering). These modifications allowed site-specific conjugation to maleimide-functionalized ASOs or siRNAs.

Using the LLQGPA–MTGase conjugation strategy, we previously observed robust Malat1 RNA knockdown in the mouse brain cortex following IV dosing only when the VHH–Fc–conjugated ASO had a 3–10–3 (16-mer) cEt full phosphorothioate chemistry (Figure 2D, E). In contrast, a 5–10–5 (20-mer) MOE-mixed backbone (MBB) ASO failed to achieve significant knockdown under comparable conditions (Figure 5A). Therefore, we were particularly interested in determining whether the alternative conjugation sites (S239C or E388C), together with Cys-Mal chemistry, could enhance the CNS activity of IV dosed 20-mer MOE-MBB ASOs, increasing the versatility of the RMT platform. Based on our goal, monovalent C5VHH–Fc constructs bearing engineered cysteine residues S239C or E388C were conjugated either to a 5–10–5 MOE-MBB Malat1 ASO carrying a 5′ maleimide group or to an Hprt siRNA bearing a 5′ maleimide modification on the sense strand (Supplementary Figure 14A, B). We next evaluated the in vivo activity of these conjugates relative to the corresponding C-terminal LLQGPA–MTGase (azide–alkyne) conjugates. qRT-PCR analysis for Malat1 RNA following IV dosing at 0.3, 1, and 3.5 mg/kg (ASO equivalents) in wild-type mice revealed that the E388C, and especially the S239C sites, combined with the Cys-Mal conjugation strategy significantly improve the potency of C5VHH-Fc-MOE-MBB Malat1 ASO in brain cortex and spinal cord compared to the C-terminal LLQGPA-MTGase approach (Figure 5A). The result is even more remarkable given the almost complete lack of activity of the latter in brain cortex (Figure 5A). In skeletal muscle (quadriceps) and heart, the three different molecules performed similarly, with the S239C Cys-Mal conjugate being overall the most potent across all the tissues examined (Figure 5A). Conjugation of Hprt siRNA to VHH-Fc via S239C slightly improved (even less so for E388C) the potency of the IV dosed molecule in the CNS of wild-type mice compared to the C-terminal LLQGPA-MTGase strategy (Figure 5B). However, when the payload is an siRNA, the C-terminal LLQGPA-MTGase conjugation method appears already efficient (Figure 2A; Figure 5B), so the differences in potencies are smaller compared to what was observed with the 20-mer MOE-MBB ASO payload (Figure 5A).

**Figure 5:**
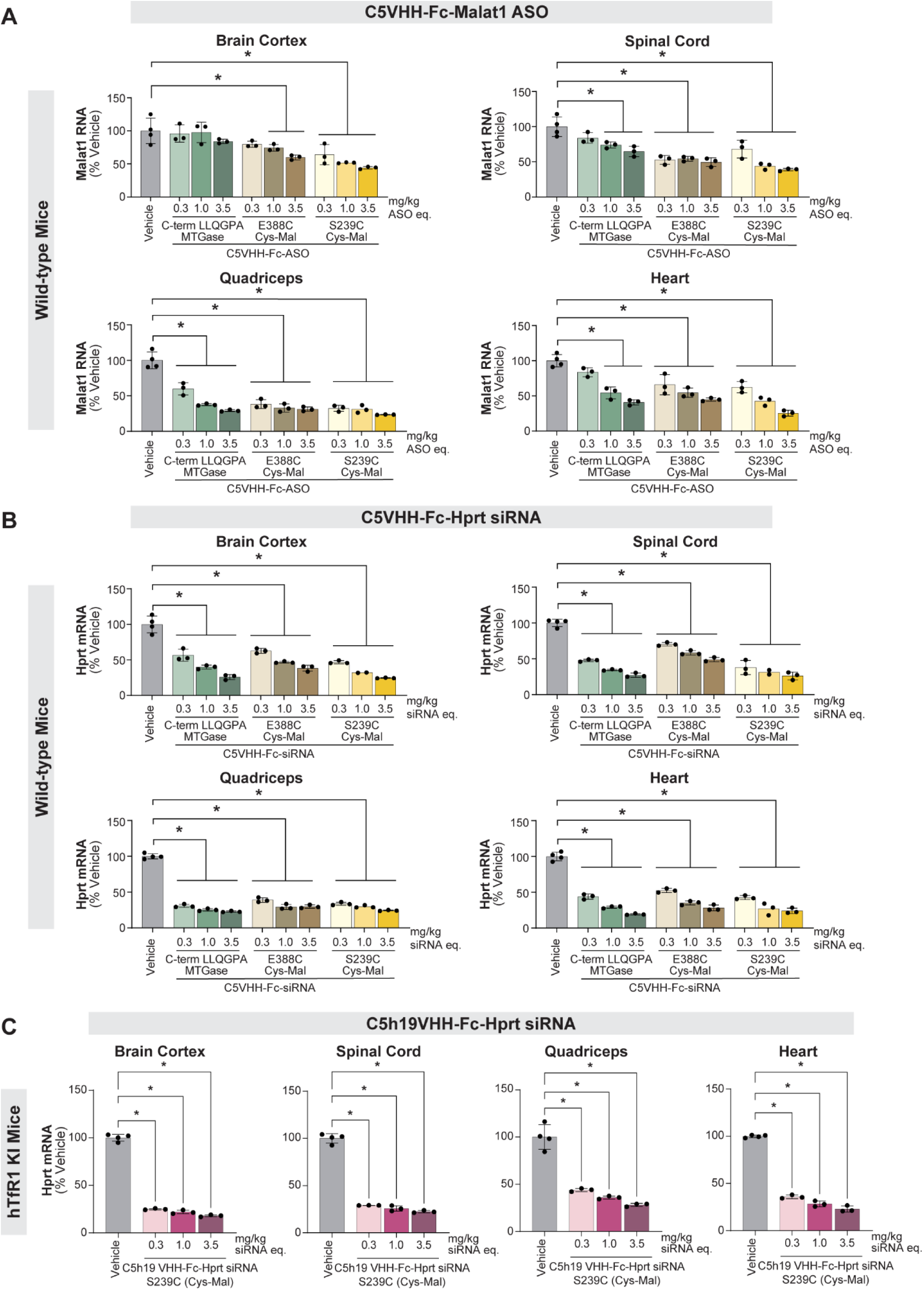
Optimization of conjugation site and chemistry improves the CNS potency of IV dosed VHH-Fc-ASO and siRNA conjugates in vivo. (**A**) Mouse Malat1 RNA levels (expressed as percentage of vehicle control animals) measured by qRT-PCR in tissues from wild-type mice dosed IV with C5VHH-Fc-Malat1 ASO (5-10-5 MOE-MBB) at the indicated dose levels (based on ASO equivalents). The ASO was conjugated using either MTGase at the LLQGPA tag (azide–alkyne), or cysteine (on Fc)–maleimide (at 5’ of ASO) (Cys-Mal) chemistry. For the latter, either E388 or S239 residues on the Fc were converted to C in order to enable the site-specific Cys-Mal conjugation. (**B**) Similarly to A, the potency of C5VHH-Fc-Hprt siRNA conjugates was assessed by qRT-PCR in different tissues from wild-type mice after IV dosing. (**C**) Mouse Hprt mRNA levels measured by qRT-PCR in tissues from human TfR1 KI (hTfR1 KI) mice dosed IV with C5h19VHH-Fc-Hprt siRNA at the indicated dose levels (based on siRNA equivalents). The siRNA was conjugated using the optimized S239C Cys (on Fc)-maleimide (at 5’ of sense strand) (Cys-Mal) strategy.

To further investigate the maximum potential of the VHH-Fc-siRNA platform, we also conjugated the maleimide-Hprt siRNA at the S239C of the C5h19-Fc fusion protein. Indeed, C5h19 is a humanized VHH clone that was identified through the human TfR1 binding affinity optimization exercise described in the previous section, and it was among the most active VHH variants when we screened the VHH-Fc-Hprt siRNA conjugates in human TfR1 KI mice (Figure 3B; Supplementary Figure 6B). The C5h19-Fc-Hprt siRNA (Cys-Mal conjugation at S239C) dosed IV in human TfR1 mice achieved exquisite potency in brain cortex, where it reached a remarkable maximum Hprt mRNA knockdown of 82% (at 3.5 mg/kg siRNA equivalents) versus vehicle control (Figure 5C). Similarly potent target reduction was measured in spinal cord, as well as in quadriceps muscle and heart (Figure 5C).

To further optimize our conjugation strategy, we explored different chemistries of the linker connecting the siRNA to the VHH-Fc at the S239C site. To that end, we introduced a single S239C mutation to C5h19VHH-Fc, and we conjugated it to the Hprt siRNA via with a variety of 5’ conjugation linkers (Supplementary Figure 15A). To directly compare Cys-Mal conjugation to SPAAC, we included maleimide linkers as well as a bromoacetamide-PEG-azide linker. The latter results in a stable thioether linkage on the VHH-Fc, and it conjugates to the BCN-siRNA (Supplementary Figure 15A). Human TfR1 transgenic mice were dosed with the different constructs IV at 0.03, 0.1, 0.3, 1.0, and 3.5 mg/kg (siRNA equivalents), a wide dose-response that enables precise assessment of the potency of the conjugates both in CNS and muscle tissues. We observed similar levels of target Hprt mRNA knockdown across the different linker constructs in the brain cortex, spinal cord, quadriceps muscle, heart, and liver (Supplementary Figure 15B), suggesting that linker chemistry is less influential than conjugation site for RMT and CNS potency of VHH-Fc-siRNA conjugates.

### Systemic dosing of an optimized C5VHH-Fc-siRNA conjugate reduces the target MAPT mRNA in NHP CNS

Equipped with the knowledge derived from our efforts around optimization of TfR1 binding affinity, VHH clone selection, and conjugation site and chemistry, we sought to prove the potential of the VHH-Fc delivery platform for RMT of siRNAs in higher primate species within the context of a therapeutically relevant target. To that end, we conjugated an siRNA targeting NHP MAPT to the C5VHH-Fc using the “S239C Cys-Mal” conjugation strategy described above. MAPT encodes the microtubule-associated protein tau, a neuronal protein that stabilizes microtubules. Tau protein becomes pathogenic when misfolded and aggregated, leading to tauopathies, such as Alzheimer’s disease and frontotemporal lobar degeneration (FTLD) (62). Therapeutic silencing of MAPT using siRNA offers a rational disease-modifying strategy by reducing total tau levels, thereby limiting tau aggregation, propagation, and downstream neurodegeneration, an approach supported by both preclinical and emerging clinical evidence (63, 64). Four weekly IV infusions of the C5VHH-Fc-MAPT siRNA in NHPs at 5 mg/kg (siRNA equivalents) resulted in more than 70% target MAPT mRNA knockdown (compared to vehicle controls) in frontal cortex, as well as temporal cortex, hippocampus, and amygdala (Figure 6). These brain regions represent key central nodes of tau accumulation and propagation in the context of tauopathies. Additional brain structures, including deeper ones, similarly show excellent target engagement by the IV dosed VHH-Fc-siRNA (Supplementary Figure 16). A notable exception is the cerebellar cortex, where the target knockdown is markedly lower relative to other brain regions (Supplementary Figure 16), similar to what was observed for the HPRT siRNA conjugate (Supplementary Figure 11). Consistent with the qRT-PCR results, quantification of tau protein by enzyme-linked immunosorbent assay (ELISA) across NHP brain regions demonstrated target protein reduction following IV administration of VHH-Fc–MAPT siRNA (Supplementary Figure 17A), with MAPT mRNA and tau protein levels showing strong concordance (Supplementary Figure 17B). The smaller reduction in tau protein relative to MAPT mRNA knockdown may reflect the long half-life of tau protein, reported to average approximately 30 days in the human CNS (65, 66). In the spinal cord, qRT-PCR revealed more than 40% target MAPT mRNA reduction compared to tissues from vehicle-dosed animals (Figure 6; Supplementary Figure 16). By virtue of TfR1 engagement, good activity was also measured in quadriceps muscle and heart of the IV dosed NHPs (Figure 6).

**Figure 6:**
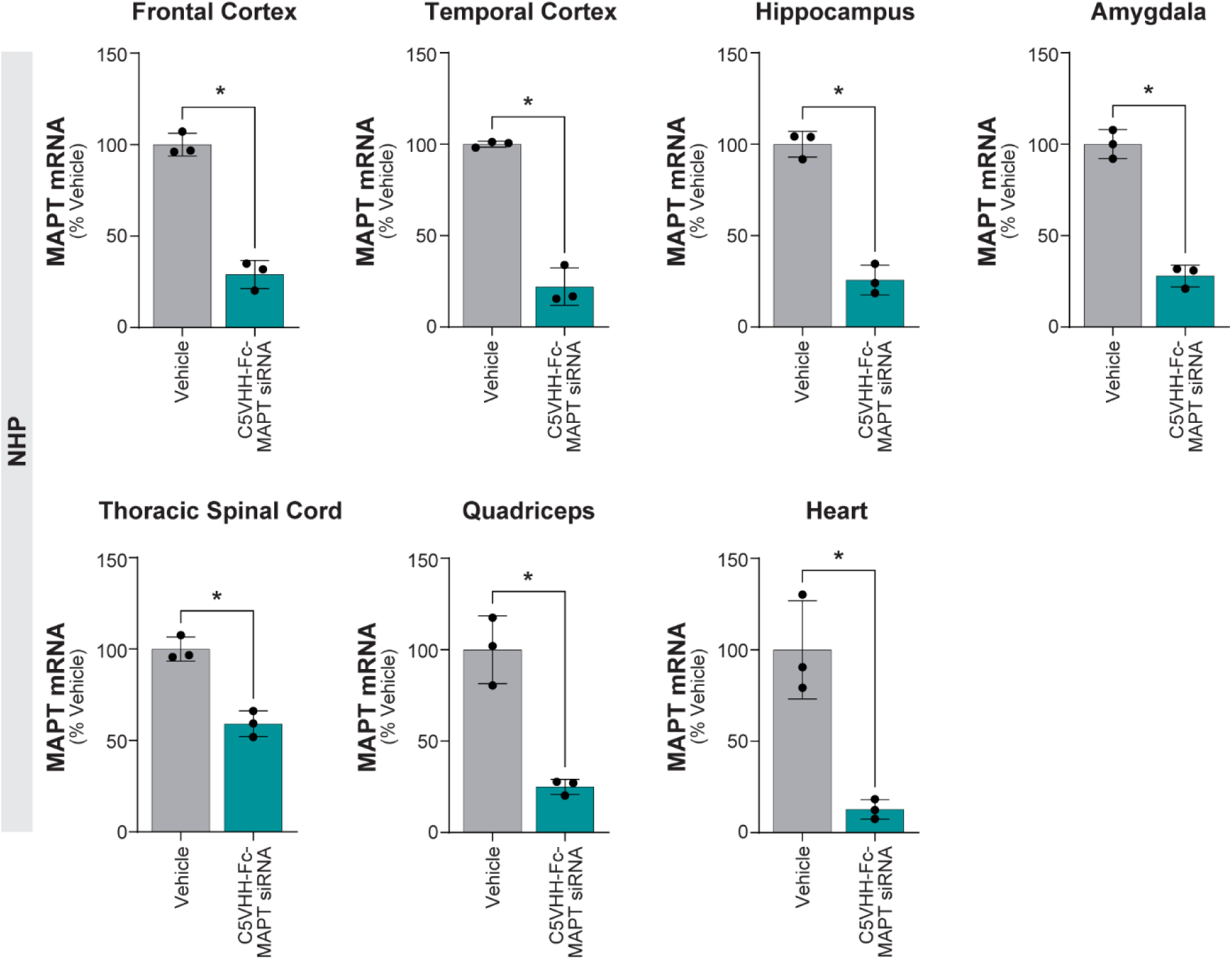
Systemic dosing of an optimized C5VHH-Fc-siRNA conjugate reduces the target MAPT mRNA in NHP CNS. MAPT mRNA levels measured by qRT-PCR in different tissues from NHPs after IV infusion with C5VHH-Fc-MAPT siRNA at 5 mg/kg (siRNA equivalents). Levels are expressed as percentage of vehicle control animals.

## DISCUSSION

In this work, we have optimized delivery of oligonucleotide cargoes across the BBB demonstrating substantial gains in activity via systematic engineering of TfR1 binding ligands. While previously published efforts utilized mAb ligands of varying complexity (28, 54), we pursued a strategy employing simplified VHH-Fc ligands that enabled us to rapidly iterate on parameters governing successful RMT and CNS target reduction, including TfR1 binding affinity and cargo conjugation site. We also built on previous work focused on ASO delivery across the BBB (28), and expanded the scope of this technology to include siRNAs, an important emerging therapeutic class for neurological indications (67–70). In so doing, we demonstrated the versatility of the VHH-Fc format for delivery of ASO and siRNA payloads to CNS tissues in rodents and NHPs, supporting potential clinical translation to humans. Consistent with the goal of enabling clinically tractable systemic administration, conjugates maintained robust CNS activity after subcutaneous dosing in NHPs, with levels of target mRNA reduction comparable to those observed after intravenous infusion.

TfR1 is a master regulator of iron homeostasis that functions largely by complexing with iron-loaded holo-Tf, internalizing for intracellular processing, and recycling to the cell surface (18). Ideal clinical TfR1 ligands should therefore avoid competition with this natural ligand for at least two reasons: achieving maximum receptor occupancy due to a lack of binding site competition, and minimum perturbation of the natural TfR1-mediated iron homeostasis. TfR1 antibodies that bind the TfR1 apical domain (AD) and avoid competition with Tf have been described (35, 36, 71), but sequence divergence between rodent and NHP TfR1 AD can limit cross-reactivity at this site. In this work, we explore a recently described family of llama-derived VHH clones that (i) binds the interface of TfR1 monomers away from natural ligand binding sites, (ii) is capable of transcytosis and functional cargo delivery into the brain of mice, and (iii) binds to mouse, rat, human, and NHP TfR1 with therapeutically relevant binding affinities, potentially simplifying clinical translation (46). Accordingly, we observed cross-species binding to TfR1 in nBRET assays and confirmed that these molecules, when expressed as monovalent Fc fusions, are taken up into human cells overexpressing mouse or human TfR1. We also employed cryo-EM to demonstrate that C5VHH family ligands indeed bind an epitope distinct from the Tf binding site. Importantly, this noncompetitive binding mode translated into no detectable perturbation of transferrin-associated iron homeostasis in vivo. Indeed, IV administration of the C5VHH-Fc-siRNA conjugate did not reduce serum iron or transferrin levels, and did not lower hemoglobin, hematocrit, red blood cell counts, or other hematological indices in mice, indicating that the conjugate does not measurably interfere with Tf/TfR1-mediated iron handling, nor causes an anemia-like phenotype under the conditions tested (Supplementary Figure 18). In previous work, mutagenesis identified human TfR1 residues E634, G636, E728, and R732 as the main functional contributors to C5VHH binding, with the epitope spanning both TfR1 monomers and being conserved across species (46). Our structure reveals a matching footprint. In our reconstruction, TfR1 E728 contacts C5VHH R40 (IMGT numbering), while TfR1 E634 engages both C5VHH Y42 and R40, and TfR1 R732 packs against C5VHH F38. TfR1 G636, also highlighted in the mutational scan, interacts with C5VHH T57 despite lacking a side chain. Beyond these previously described contacts, our cryo-EM map resolves additional stabilizing interactions not captured in the mutagenesis dataset: TfR1 R235 forms a salt bridge with C5VHH D29, TfR1 Y309 also engages C5VHH D29, and TfR1 P323 packs against C5VHH L107 and F38, contributing to conformational complementarity. We also observe that TfR1 NAG802 is accommodated by C5VHH N64 and T59, suggesting a glycan-supported stabilization of the paratope–epitope interface. Together, these structural observations provide direct experimental confirmation of the binding mode proposed by prior epitope mapping studies and extend it by defining distinct contacts formed in the C5VHH-TfR1 complex.

High binding affinity for TfR1 negatively correlates with BBB transcytosis of ligands (28, 34, 36, 72) but supports distribution to peripheral tissues (5, 8, 9). Our observations agree with these reports. Initial testing of C5VHH-Fc conjugates in wild-type mice, where binding affinity is relatively lower, resulted in greater than 50% target reduction in brain cortex following two IV administrations at 1 mg/kg siRNA or ASO equivalents. When targeting Hprt, we demonstrated concomitant protein reduction throughout the brain indicating that knockdown was not restricted to brain vasculature. Upon testing these conjugates in transgenic mice bearing human TfR1, where binding affinity is sub-nM, we observed substantially lower activity in CNS tissues, with target reduction in brain cortex failing to surpass 50% after two IV administrations at 1 mg/kg. In contrast, target reduction in quadriceps muscle and heart was roughly comparable between mouse strains. We hypothesized that variants with lower binding affinities than C5VHH-Fc would be more active in human TfR1 transgenic mice. Indeed, when we administered siRNA conjugates of humanized C5VHH-Fc derivatives with a broad range of binding affinities to wild-type and human TfR1 transgenic mice, we observed the greatest target reduction in brain cortex within an intermediate range of TfR1 binding affinities (nBRET Ki approximately 10-100 nM). We also repeated the observation that knockdown in quadriceps muscle and heart is independent of binding affinities in the tested range under our dosing paradigm. Testing the closely related, but distinct, B8VHH family yielded similar results, supporting the notion that the most active conjugates in rodents possess TfR1 binding affinities in the 10-100 nM range. Of note, we tested in NHP four conjugates representing a similar range of binding affinities, and observed the greatest target reduction with C5VHH-Fc-siHPRT, which has a nBRET affinity to NHP TfR1 of 10 nM. In NHPs, the observed drop-off in activity with lower binding affinity conjugates was sharper than in rodents. We do not rule out the possibility that sequence differences between the C5VHH and B8VHH families affect binding kinetics and pharmacokinetic properties of conjugates, rendering the B8VHH family less potent in NHP. Notably, in both mice and NHPs we also observed substantial target mRNA reduction in dorsal root ganglia and peripheral nerves, including trigeminal and sciatic nerve, after systemic dosing, indicating that the same VHH-Fc conjugates can engage targets across the blood–nerve barrier in addition to the BBB.

A potential strategy to boost the activity of BBB-crossing conjugates is to increase the number of cargoes transported by a single receptor through increased DAR, but it has been reported that DAR higher than 1 is detrimental to the pharmacokinetics and activity of antibody-siRNA conjugates (73) in skeletal muscle. Because these studies omit delivery across the BBB, we sought to close this knowledge gap. We generated DAR1 and DAR2 VHH-Fc-siRNA conjugates and compared their activity in the CNS and in systemic tissues of mice. We observed overall less target reduction in CNS tissues of animals treated with DAR2 conjugates at all dose levels. A similar, albeit weaker, trend was observed in skeletal and cardiac muscle; however, the minimal differences across dose levels suggest saturation. On the other hand, target reduction in the liver of animals dosed with DAR2 conjugates was greater than in animals dosed with DAR1 conjugates, suggesting that higher DAR leads to increased uptake and activity in the liver, consistent with published results (73).

Conjugation site has been shown to influence activity, stability, pharmacokinetics, and tolerability of ADCs (52, 58, 59). Optimization of the conjugation site has also been shown to improve the PK properties of antibody–oligonucleotide conjugates (AOCs) (54), but to our knowledge no published work has examined the impact of conjugation site on siRNA conjugate activity across the BBB. We tested siRNA and ASO (5-10-5 MOE-MBB gapmer) conjugated to VHH-Fc at three different positions: C-terminal, position 239 (CH2 domain), and position 388 (CH3 domain). For siRNA conjugates, we observed slight differences in potency in brain cortex by conjugation site, with the greatest target knockdown measured in animals treated with the position 239 conjugate. For the ASO conjugates, we observed little target reduction in CNS tissues with C-terminal conjugates, while conjugation to positions 239 and 388 yielded more substantial reduction. Position 239 conjugate achieved target reductions surpassing 50% in brain cortex and spinal cord. Interestingly, the potency in quadriceps muscle and heart was less affected by conjugation site, though similar trends to CNS tissues emerged, with position 239 conjugation being superior for siRNA and ASO across all tissues. Although elucidating the mechanisms underlying this improvement is beyond the scope of the present study, future work could investigate whether conjugation at position 239 enhances Fc-mediated shielding of ASO phosphorothioate groups, thereby improving pharmacokinetics and/or transcytosis relative to conjugation at the other sites. Indeed, it was recently demonstrated that shielding ASO PS groups by means of a PS-ASO binding Fab fragment improved plasma PK, as well as brain exposure of AOCs (74). The robust CNS activity achieved with different antisense modalities (i.e., ASO, siRNA) and chemical modifications (i.e., cEt, MOE-MBB) via optimized conjugation site and chemistry indicates that our VHH-Fc platform aligns with the overarching goal of developing a versatile delivery system for diverse therapeutic oligonucleotides.

Optimization of linker chemistry is critical for cytotoxic ADCs (75–78), but previous work with AOCs suggests that potency and PK are minimally affected by this parameter when controlling for conjugation site (54, 73). We investigated a series of thiol reactive linkers at our most active conjugation site and observed nearly identical potency among them. The factors considered in our structure–activity relationship (SAR) analysis were the role of the phosphorus linkage between oligo and ligand (C_3_ maleimide plus PO or PS), the importance of preventing retro-Michael deconjugation (C_3_ maleimide closed, C_3_ maleimide open, SMCC), the stability of the bond between the hexylamino-oligonucleotide and linker, specifically carbamate linkages (maleimide-PEG-carbamate) and amide linkages (SMCC and C_3_ maleimide), the presence of a short PEG linker (maleimide-PEG_1_-carbamate and acetamide-PEG_2_-azide-BCN), and the difference between maleimide and SPAAC (acetamide-PEG_2_-azide-BCN). We observed nearly identical target reduction with all linkers tested, both in CNS and muscle tissues, in agreement with previously published work on AOC linker chemistry (73). Future studies may also examine linkers engineered for specific release by enzymes including cathepsins (Val-Cit-PABC linkers), β-glucuronidase (β-glucuronide linkers), or other cell and tissue type specific enzymes, although previous work suggests there is no benefit in the case of Val-Cit (73).

Finally, we demonstrated robust reduction of a therapeutically relevant target, MAPT, across multiple brain regions in NHP after treatment with C5VHH-Fc-MAPT siRNA. Demonstrating MAPT suppression across frontal and temporal cortical regions, hippocampus, and amygdala in NHP provides a biologically and translationally relevant framework for evaluating BBB-delivered tau-lowering therapeutics in humans. These results represent the culmination of our systematic optimization of the VHH-Fc–based RMT platform, including selection of TfR1 ligands with appropriate binding affinities and engineering of conjugation site and chemistry to support efficient transport and activity of both siRNA and ASO payloads. The ability of VHH-Fc conjugates to drive robust target engagement in the CNS after intravenous and subcutaneous dosing highlights the potential of this RMT platform to serve as a basis for the identification of future systemically delivered antisense therapeutics for neurological diseases.

## Supporting information

Huggins, Carrer et al. 2026_Supplementary Data

## ABBREVIATIONS

ASO: antisense oligonucleotide
BBB: blood-brain barrier
BRET: bioluminescence resonance energy transfer
cEt: 2′-constrained ethyl
CNS: central nervous system
Cys-Mal: cysteine–maleimide
DAR: drug to antibody ratio
eq: equivalents
Fc: fragment crystallizable region
Hprt: hypoxanthine phosphoribosyltransferase 1
IV: intravenous
Kd: equilibrium binding affinity
Ki: inhibition constants
KI: knock-in
Malat1: metastasis-associated lung adenocarcinoma transcript 1
MAPT: microtubule-associated protein tau
MBB: mixed backbone
MOE: 2′-O-methoxyethyl
MTGase: microbial transglutaminase
nBRET: nanoBRET
NHP: non-human primate
PO: phosphodiester
PS: phosphorothioate
RMT: receptor-mediated transcytosis
SC: subcutaneous
siRNA: small interfering RNA
Tf: transferrin
TfR1: transferrin receptor 1
VHH: variable heavy domain of heavy chain-only antibody

## ACKNOWLEDGMENTS

We thank Alena Delio and Jamie Watson for histological support; Kaitlyn Trinh, Faye Lopez, Nicole Lombardo, and Susan Li for technical support with animal dosing, tissue collection and processing, and ELISA assays; Tracy Reigle and Raul Alonzo for assistance with graphics and figure preparation.

## DATA AVAILABILITY

All data are available upon request.

