## Supplementary material for "Delivery of small interfering RNA and antisense oligonucleotides across the blood-brain barrier with monovalent transferrin receptor 1 binding VHH-Fc fusion proteins": Huggins, Carrer et al. 2026_Supplementary Data

Author contributions: IJH, MC, JAS, MF, BH, SP, TPP, MA, MAB, SKK, RG-M, AAR, FK, HG, AC, MB-H, AP-D, and RQ conducted experiments and analyzed the data. IJH, MC, GJ, MD, FR, HBK, HTZ, PJ-n, MT, and EES supervised the studies. IJH and MC wrote the paper with input from all authors.

### SUPPLEMENTARY MATERIALS AND METHODS

**Synthesis of Malat1 ASO containing 5’-SMCC linker**

A 2-dram vial equipped with a magnetic stir bar was charged with 5′-aminohexyl oligonucleotide (200 mg, 25.7 µmol) and 0.2 M phosphate buffer (pH 7.5, 1.9 mL). The mixture was stirred at room temperature until a homogeneous solution was obtained. In a separate vial, 2,5-dioxopyrrolidin-1-yl 4-(2,5-dioxo-2,5-dihydro-1H-pyrrol-1-yl)benzoate (SMCC NHS ester, 15 mg, 45.0 µmol) was dissolved in acetonitrile (1.5 mL), and the resulting solution was added to the oligonucleotide solution. The reaction mixture was stirred at room temperature for 22 hours and then heated to 35°C for 3 hours. Completion of the reaction was confirmed by LC–MS analysis. To remove excess unreacted linker, the oligonucleotide product was precipitated by the addition of 4 volumes of ethanol. After the precipitate had settled, the supernatant was decanted, and the solid was reconstituted in <5 mL of 0.2 M sodium acetate solution. The oligonucleotide was then reprecipitated with 4 volumes of ethanol. After the precipitate had settled, the supernatant was decanted, and the solid was reconstituted in <5 mL of deionized water. The resulting solution was lyophilized to afford the product as a white solid.

**Synthesis of HPRT siRNA sense strand containing 5′-SMCC linker**

To a solution of 5′-hexylamino oligonucleotide (142 mg, 19.97 µmol) in 0.05 M sodium phosphate buffer (pH 7.0, 1.5 mL) was added a solution of SMCC NHS ester (63.0 mg, 188.5 µmol) in DMSO (1 mL). The reaction mixture was stirred at room temperature for 18 hours, then diluted with water (40 mL) and purified by SAX HPLC using Source 30Q resin (Cytiva Cat. 17-1275-03); eluent A: 100 mM ammonium acetate in 30% aqueous acetonitrile; eluent B: 1.5 M NaBr in A; gradient: 0–60% B over 60 minutes; flow rate: 14 mL per minute. Fractions containing the full-length oligonucleotide, as determined by LC–MS analysis, were combined and concentrated under reduced pressure. The resulting residue was dissolved in water and desalted by reverse-phase high-performance liquid chromatography (HPLC) to afford the desired product (117 mg, 80%).

**Synthesis of HPRT siRNA sense strand containing 5'-maleimide-ethoxyethyl-carbamate linker**

To a solution of 1-(2-(2-hydroxyethoxy)ethyl)-1H-pyrrole-2,5-dione ($PEG_1$-maleimide, 100 mg, 540 µmol) in anhydrous DMF (0.5 mL) were added di(*N*-succinimidyl) carbonate (140 mg, 540 µmol) and triethylamine (0.1 mL, 810 µmol). The resulting slurry was vortexed for 5 minutes until a clear solution was obtained. After stirring at room temperature for 1 hour, 0.1 M HCl (0.01 mL, 0.81 mmol) was added to yield the activated maleimide-$PEG_1$-NHS carbonate linker. Separately, 5′-hexylamino oligonucleotide (80 mg, 11.25 µmol) was dissolved in 0.05 M sodium phosphate buffer (pH 7.0, 1 mL). A portion of the activated linker (104 µL, 112.5 µmol) was then added. The reaction mixture was stirred at room temperature for 1 hour, diluted with water (40 mL), and purified and desalted by HPLC using the same procedure described for the synthesis of SMCC HPRT sense strand to yield the desired product (62 mg, 75%).

**Cryo-electron microscopy and structure determination of TfR1:Tf:C5VHH complexes**

Cryo-EM sample preparation, data collection, data processing, and model building were performed by NanoImaging Services, Inc. as follows:

*Cryo-EM grid preparation and data acquisition*

The TfR-Tf-C5VHH complex was assembled by mixing recombinant human transferrin receptor 1 (TfR1) (ACROBiosystems, Cat. TFR-H5213), purified transferrin (Tf) from human serum (Jackson ImmunoResearch, Cat. 009-000-050), and C5VHH-6xHis (produced at Ionis) at a molar ratio of 1:2:4 and incubating for 1 hour at room temperature. 3.0 µL of the complex was applied to graphene oxide coated UltrAuFoil R 1.2/1.3 grids (Quantifoil, Cat. Q350AR13A) that had been glow discharged with a PELCO easiGlow™ instrument (Ted Pella, Cat. 91000). Excess sample was blotted and grids were plunge-frozen in liquid ethane using Vitrobot Mark IV (Thermo Fisher Scientific). Data collection was carried out using a FEI Titan Krios (Thermo Fisher Scientific) transmission electron microscope operated at 300kV and equipped with a Quantum 967 LS imaging filter (Gatan) and K3 Direct Detection Camera (Gatan). A total of 7,051 high magnification images were collected using Leginon software (1) at 10,5000x magnification with a calibrated pixel size of 0.83 Å/ pixel. A total dose of 50.1 $e^{-}/Å^2$ was applied over 35 frames with a total exposure time of 1.4 s. Data were collected at a nominal defocus range of −0.8 to −1.8 µm.

*Cryo-EM data processing*

All data processing was carried out in cryoSPARC 4.6 (2). Images were preprocessed on-the-fly with CryoSPARC Live 4.6, and 5,729 micrographs were selected based on a CTF fit cutoff of 4 Å. 5.3 million particles were picked using a template picker and extracted. The extracted particles were subjected to three rounds of 2D classification and the best particles representing the intact complex were used to generate five ab initio models. Several iterative rounds of heterogeneous refinement, homogenous refinement, and non-uniform refinement were performed to select the homogenous conformation for final reconstruction. The final 3D reconstruction (C1 symmetry imposed) was generated using 418,000 particles and has a nominal resolution of 2.46 Å.

*Model building*

The TfR-Tf complex structure (PDB: 1SUV) and VHH (PDB: 6DBA) were docked into the cryo-EM map and used as an initial template for model building. Iterative rounds of manual adjustment in COOT (3) and real-space refinement in Phenix (4) were performed against the cryo-EM map to improve the stereochemistry as well as the model map coefficient correlation. Validation of the final model was performed using MolProbity in Phenix. For image generation, density maps were rendered in UCSF-ChimeraX version 1.9 (2024-12-11) (5) with a contour level of 0.267. Detailed structural images were made using Discovery Studio Simulation Client version 26.2.100 (Dassault Systèmes, BIOVIA).

**Histological analysis**

Tissue specimens from brain of mice and NHP were immersed in 10% neutral buffered formalin for 48–72 hours. Following paraffin embedding, sections were cut at a thickness of 4 µm and mounted onto glass slides. Immunohistochemical staining was carried out using a Ventana Discovery Ultra automated staining platform. HPRT protein was detected using a rabbit anti-HPRT monoclonal antibody (Abcam, Cat.

Ab109021) on a Ventana Ultra staining system. Slides were treated with heat induced antigen retrieval (HIER) with Ventana CC1 solution (Ventana, Cat. 950-500) for 64 minutes. The tissue was permeabilized with 5% Tween 20 (VWR, Cat. 0777-1L) for 32 minutes at 37°C. The primary antibody was diluted with Discovery Antibody Diluent with casein (Ventana, Cat. 760-219) at 1:800, then incubated for 1 hour at 37°C. The antibody was detected with OmniMap anti-rabbit polymer detection (Ventana, Cat. 760-4311). The rabbit polymer was labeled with Ventana ChromoMap DAB kit (Ventana, Cat. 760-159). Whole-slide images were acquired using a Hamamatsu S360 digital slide scanner at 20x magnification. Histological staining intensity and distribution were evaluated by a board-certified pathologist using light microscopy and assigned semi-quantitative scores.

**Measurement of protein levels in non-human primate tissues**

Total tau protein levels were quantified in non-human primate (NHP) frontal cortex homogenates using the Human Total Tau ELISA Kit (Thermo Fisher Scientific, Cat. KHB0041) according to the manufacturer's instructions. Tissue punches from cortical regions of interest were homogenized in RIPA buffer supplemented with a protease and phosphatase inhibitor cocktail (Thermo Fisher Scientific, Cat. 1861281) and EDTA (Thermo Fisher Scientific, Cat. 1861274). Lysates were clarified by centrifugation at 5,000 g for 10 minutes at 4°C, and the clarified supernatant was collected for analysis. Total protein concentration was determined separately for each clarified lysate and used for normalization. ELISA standards and clarified tissue lysates were assayed in duplicate according to the kit protocol. For occipital cortex, samples were assayed at multiple dilutions, and the 1:4,000 dilution (with an additional 1:2 dilution in well) was used for analysis because the measured absorbance values fell within the quantifiable range of the 4-parameter logistic standard. For the remaining brain regions, samples were assayed on a separate plate with approximately the same amount of total protein loaded per well. Following incubation and wash steps, bound tau was detected using the detection reagents and substrate solution provided with the kit. Absorbance was measured using a SpectraMax Plus 384 microplate reader (Molecular Devices) at 450 nm, and total tau concentrations were interpolated from the standard curve. Values were corrected for dilution where applicable, normalized to total protein concentration for each sample, and expressed as percent change relative to the corresponding control group.

HPRT protein levels were quantified in NHP brain tissue homogenates using the Simple Western automated Western blot system (Bio-Techne). 4-mm tissue punches from brain cortex and caudate were homogenized in RIPA buffer supplemented with a protease and phosphatase inhibitor cocktail (Thermo Fisher Scientific, Cat. 1861281). Lysates were clarified by centrifugation at 5,000 g for 10 minutes at 4°C, and the clarified supernatants were collected for analysis. Total protein concentration was determined using Pierce Rapid Gold BCA Protein Assay Kit (Thermo Fisher Scientific, Cat. A53225) separately for each clarified lysate and used for normalization. Primary antibody incubations were performed with a rabbit polyclonal anti-HPRT antibody (Proteintech, Cat. 15059-1-AP) diluted 1:1,000 in antibody diluent buffer (Bio-Techne, Cat. 042-203) and a rabbit monoclonal anti-GAPDH antibody (14C10; Cell Signaling Technology, Cat. 2118) diluted 1:1,000 in the same buffer. Signal was developed using the ProteinSimple Anti-Rabbit Detection Module (Bio-Techne, Cat. DM-001). The resulting signal intensities were quantified using Compass for Simple Western software (Bio-Techne). HPRT signal was normalized to GAPDH as a loading control, and HPRT protein levels were expressed as percent change relative to vehicle-treated control animals.

**Hematology and serum iron, transferrin measurements in mice**

Adult female wild-type C57BL/6NTac mice were dosed by intravenous (IV) injection with either vehicle (PBS) or C5VHH-Fc-Hprt siRNA at 1, 3.5, 5, or 10 mg/kg (siRNA equivalents) on study days 1 and 8. The mice were euthanized on day 22. Whole blood was collected at the end of the study by cardiac puncture under terminal anesthesia and placed in Microvette™ 500 EDTA K2E tubes (SARSTEDT, Cat. 20.1339.100). Hematology indices were determined from complete blood count (CBC) measurements performed at IDEXX Laboratories. Serum iron and transferrin values in mice were also assessed at study termination. Whole blood was collected by cardiac puncture under terminal anesthesia, and serum was separated using MiniCollect™ capillary blood collection system tubes (Greiner Bio-One, Cat. 22030401).

The tubes were kept at room temperature, then spun at 3,000 g in an Eppendorf benchtop microcentrifuge at room temperature for 10 minutes. The separated serum was then transferred to a 96-well plate and stored at -20°C until further analysis. Iron and transferrin levels in the serum were measured using a Beckman Coulter AU480 Chemistry Analyzer (Beckman Coulter). All mouse procedures complied with the eighth edition of the Guide for the Care and Use of Laboratory Animals (AAALAC-accredited Unit 000962 and State of California Department of Public Health Certificate No.071) and were conducted under the Institutional Animal Care and Use Committee (IACUC) protocol 2023-1219.

### SUPPLEMENTARY TABLES AND FIGURES

| Gene | Species | Forward Primer | Reverse Primer | Probe | | |
|---|---|---|---|---|---|---|
| | | Sequence | Sequence | Sequence | Fluorescent Reporter/Label | Quencher(s) |
| Hprt1 | Mouse | 5'-CTCCTCAGACCGCTTTTTGC-3' | 5'-TAACCTGGTTCATCATCGCTAATC-3' | 5'-CCGTCATGCCGACCCGCAGT-3' | 5' FAM | Internal ZEN™/ 3'-Iowa Black™ FQ |
| Malat1 | Mouse | 5'-TGGGTTAGAGAAGGCGTGTACTG-3' | 5'-TCAGCGGCAACTGGGAAA-3' | 5'-CGTTGGCACGACACCTTCAGGGACT-3' | 5' FAM | Internal ZEN™/ 3'-Iowa Black™ FQ |
| Gapdh | Mouse | 5'-GGCAAATTCAACGGCACAGT-3' | 5'-GGGTCTCGCTCCTGGAAGAT-3' | 5'-AAGGCCGAGAATGGGAAGCTTGTCATC-3' | 5' FAM | Internal ZEN™/ 3'-Iowa Black™ FQ |
| HPRT1 | *Macaca fascicularis* (NHP) | Taqman™ Gene Expression Assay from Thermo Fisher Scientific (Assay ID: Rh02800695_m1) | | | 5' FAM | 3'-MGB NFQ |
| MAPT | *Macaca fascicularis* (NHP) | 5'-AGGACAGAGTGCAGTCGAAGATC-3' | 5'-AGGTCAGCTTGTGGGTTTCAA-3' | 5'-CACCCATGTCCCTGGCGGAGG-3' | 5' FAM | Internal ZEN™/ 3'-Iowa Black™ FQ |
| GAPDH | *Macaca fascicularis* (NHP) | 5'-CAACGGATTTGGTCGTATTGG-3' | 5'-GGCAACAATATCCACTTTACCAGAGT-3' | 5'-CGCCTGGTCACCAGGGCTGCT-3' | 5' FAM | Internal ZEN™/ 3'-Iowa Black™ FQ |

**Supplementary Table 1:** Sequences of primer-probe sets used in qRT-PCR assays. Internal ZEN/3'-Iowa Black™ FQ is a double-quencher system. MGB: minor groove binder. NFQ: nonfluorescent quencher.

**nBRET Binding Affinity Measurement**
Ki (nM) for TfR1 from Various Species Measured at pH 7.4

| TfR1 Species → | Human TfR1 | | *Macaca fascicularis* (NHP) TfR1 | | Mouse TfR1 |
|---|---|---|---|---|---|
| VHH Clone ↓ | VHH-Fc | VHH-Fc-Hprt siRNA | VHH-Fc | VHH-Fc-HPRT siRNA | VHH-Fc-Hprt siRNA |
| B8V12 | 2.0 | 1.5 | 37.1 | 7.2 | 534.8 |
| B8V16 | 13.4 | 22.8 | 1091.0 | 630.1 | 15.3 |
| B8V27 | 4.0 | 7.0 | 100.1 | 45.9 | 269.1 |
| B8V31h4 | 103.8 | 535.5 | 641.6 | 366.2 | 2588.0 |
| B8V32h14 | - | 58.2 | - | 74.1 | 6755.0 |
| B8V35 | 18.5 | 40.5 | 90.8 | 57.3 | 4108.0 |
| B8V37 | 30.6 | 202.1 | 1198.0 | 1373.0 | NBD |
| B8V40 | 41.8 | 41.4 | 113.6 | 43.5 | 2157.0 |
| C5 | 0.8 | 0.4 | 11.9 | 9.9 | 24.5 |
| C5h19 | 6.0 | 7.8 | 1614.0 | 612.5 | 958.3 |
| C5h20 | 4.5 | 2.3 | 445.8 | 90.4 | 9.4 |
| C5h9 | 6.0 | 18.5 | 1091.0 | 1757.0 | 260.0 |
| C5V30 | 44.3 | 150.9 | 1297.0 | 437.4 | 1726.0 |

**Supplementary Table 2:** Binding affinity (Ki) of different VHH-Fc fusion proteins, and VHH-Fc-siRNA conjugates for TfR1 in various species, as measured by nanoBRET (nBRET) assay at pH7.4. The various VHH-Fc fusion proteins and VHH-Fc-siRNA conjugates contain different VHH clones from both C5- and B8- families. The VHH portion of the molecule represents the TfR1 ligand, so the identity of the VHH clone is specified in the left column.

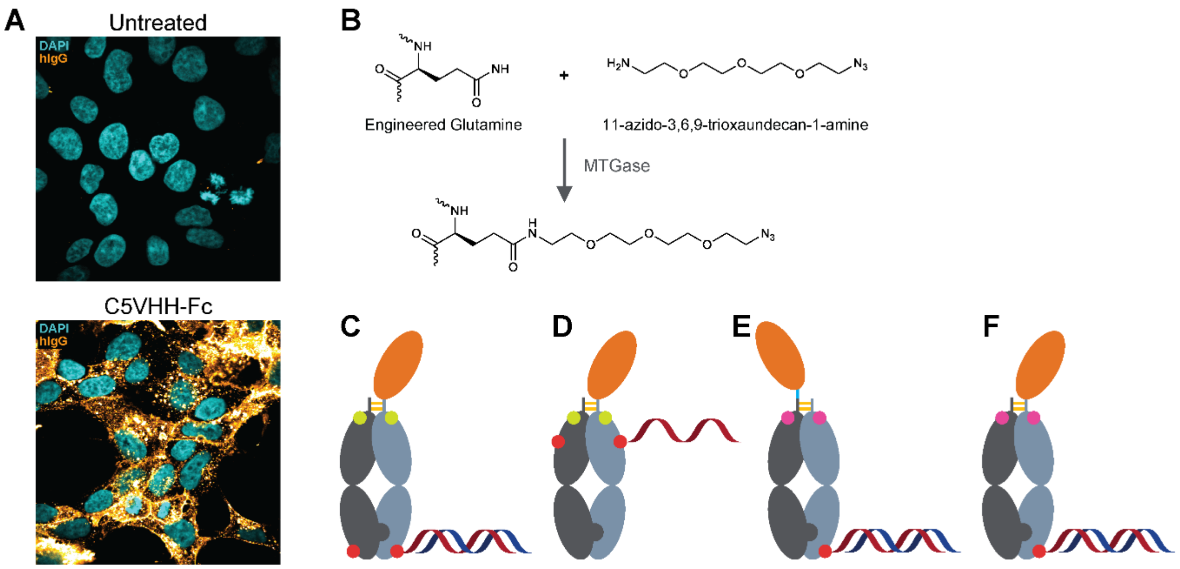


**Supplementary Figure 1:** Human TfR1-binding VHH molecules are efficiently taken up into HEK293 cells that overexpress human TfR1. (**A**) Confocal micrographs of HEK293 cells expressing human TfR1, either untreated or treated with C5VHH-Fc, and stained with DAPI (cyan) and anti-hIgG-FITC (gold). (**B**) Schematic representation of the enzymatic reaction used to label monovalent VHH-Fc proteins with a bifunctional amino-PEG$_3$-azide linker via microbial transglutaminase (MTGase). (**C-E**) Schematic illustrations of (**C, F**) VHH-Fc-conjugated siRNA via C-terminal transglutaminase LLQGPA tag, (**D**) VHH-Fc-conjugated ASO via internal LLQG tag (inserted at Fc positions 294-297), or (**E**) VHH-Fc-conjugated siRNA via C-terminal Myc tag (EQKLISEEDL). The "knobs-into-holes" heterodimeric configuration is shown with the VHH ligand (orange) fused to the Fc (grey). T366W-bearing "knob" chains are dark grey while T366S, L368A, Y407V-bearing "hole" chains are light grey. Immune effector function modulating modifications are represented either by a green dot ("LALA": L234A, L235A) or by a pink dot ("LALAPG": L234A, L235A, P329G). Conjugation sites are indicated by red dots. Ligands with two conjugation sites (C and D) are shown with one oligonucleotide randomly attached at either site (DAR1); however, both sites are marked with red dots to indicate the possible DAR2 conjugates. Hinge disulfide bonds are represented by gold bars. The siRNA and ASO are shown as duplexed and single stands, respectively. A flexible "GGGGSGGGGS" linker between VHH and Fc is denoted by a blue line segment in E.

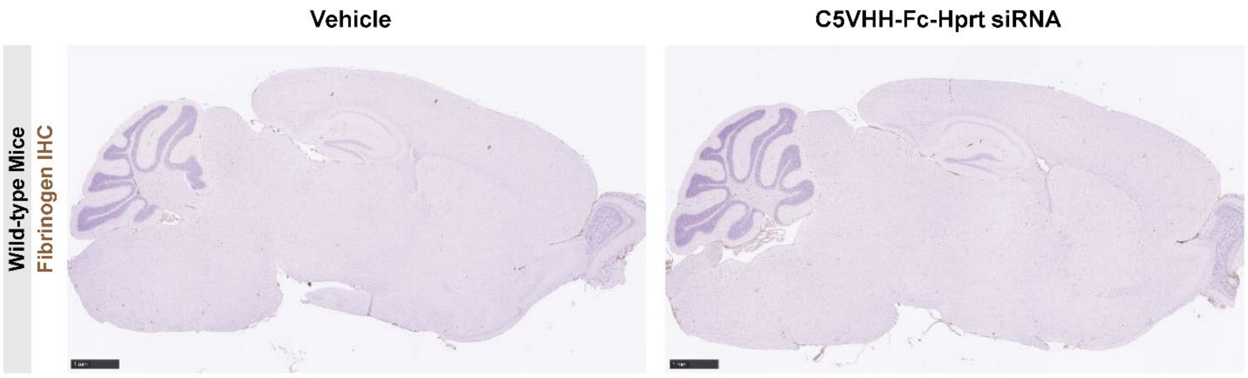


**Supplementary Figure 2:** C5VHH-Fc-Hprt siRNA does not cause blood-brain barrier (BBB) damage. Fibrinogen immunohistochemistry (IHC) performed on histological sections of brain from mice dosed via IV injection with either vehicle (PBS) or 1 mg/kg (siRNA equivalents) of C5VHH-Fc-Hprt siRNA on study days 1, 8. Mice were euthanized, and tissue collected, on day 22. Scale bars: 1 mm.

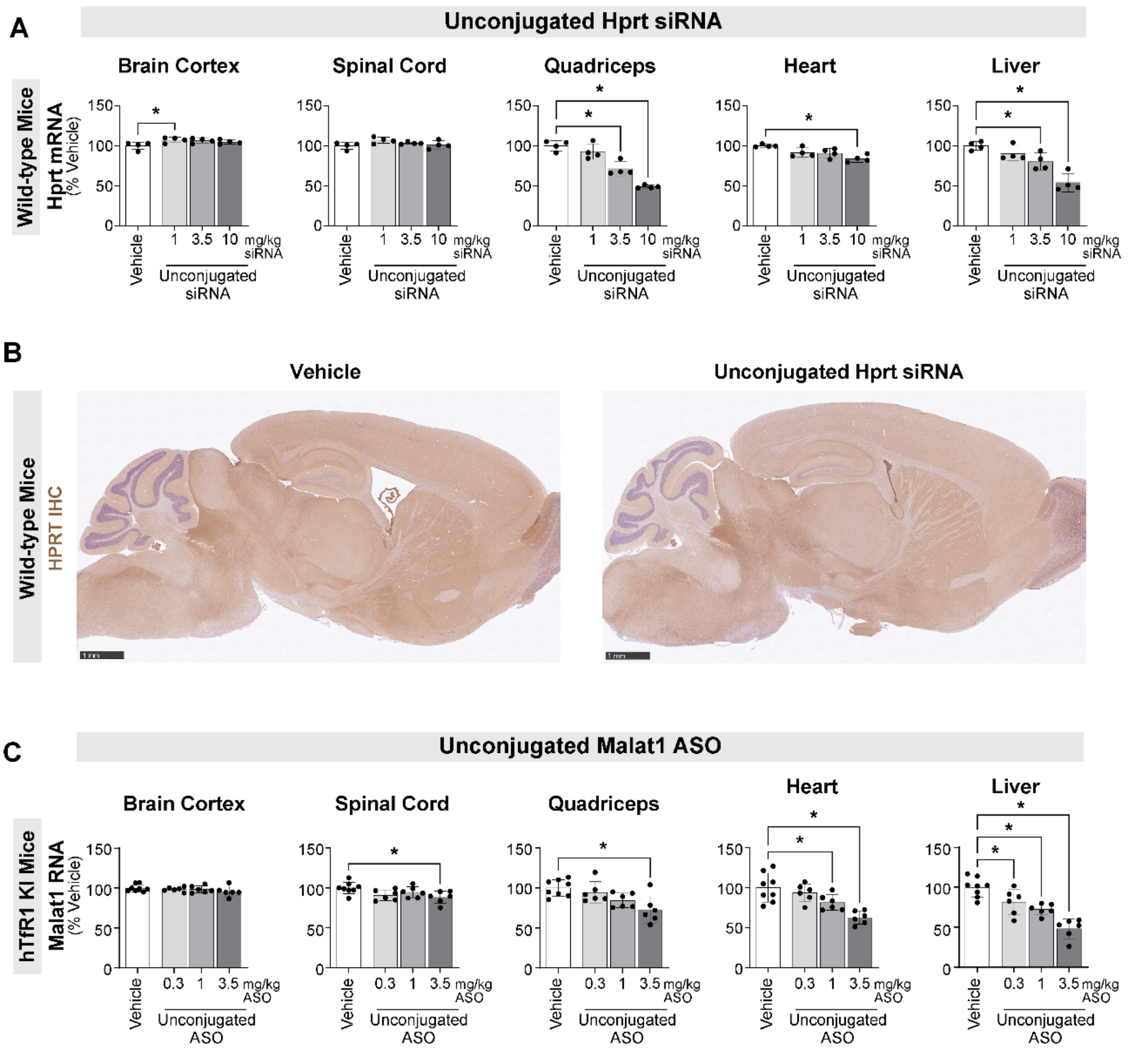


**Supplementary Figure 3:** Unconjugated Hprt siRNA and Malat1 ASO dosed IV are not active in the murine CNS. (**A**) Target Hprt mRNA knockdown measured by qRT-PCR in key CNS and systemic tissues from wild-type mice dosed IV with unconjugated Hprt siRNA on study days 1 and 8 at 1, 3.5, and 10 mg/kg. The mice were euthanized on study day 22. No activity was observed in CNS, while heart and especially quadriceps muscle and liver show good activity, therefore serving as internal controls for the validity of the experiment. (**B**) Immunohistochemistry (IHC) to detect HPRT protein (brown stain) in histological sections of brain from wild-type mice dosed IV with unconjugated Hprt siRNA at 1 mg/kg. Scale bars: 1 mm. (**C**) Target Malat1 RNA knockdown measured by qRT-PCR in key CNS and systemic tissues from human TfR1 KI (hTfR1 KI) mice dosed IV with unconjugated full PS 3-10-3 cEt gapmer Malat1 ASO on study days 1, 8, and 15 at 0.3, 1, and 3.5 mg/kg. The mice were euthanized on study day 22.

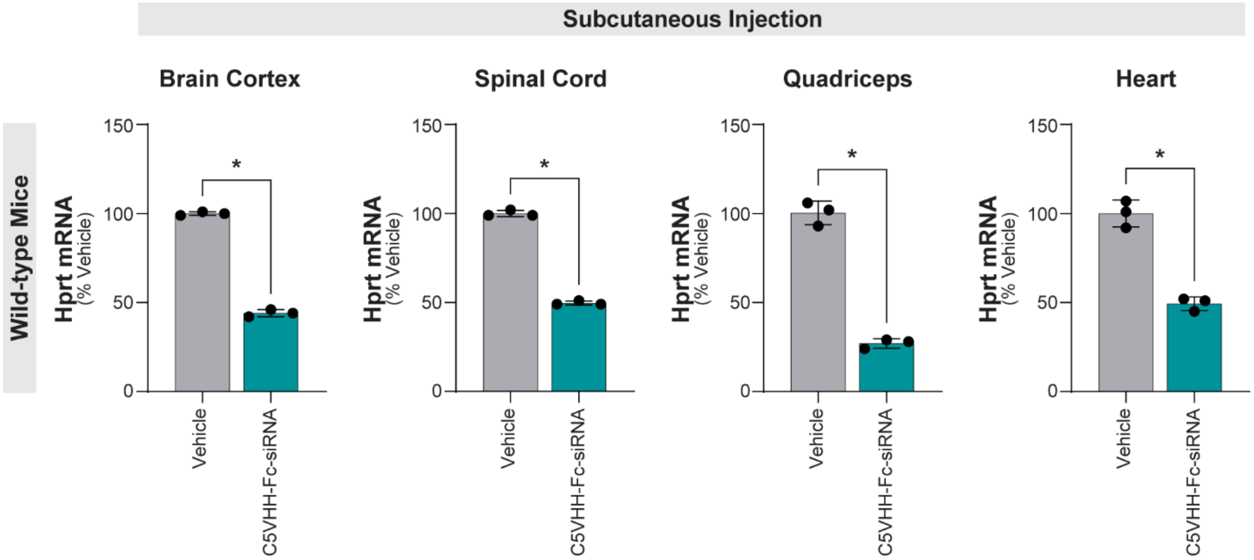


**Supplementary Figure 4:** Subcutaneous delivery of C5VHH-Fc-siRNA results in robust CNS activity in mice. Target Hprt mRNA knockdown measured by qRT-PCR in key CNS and systemic tissues from wild-type mice dosed via subcutaneous injection with C5VHH-Fc-Hprt siRNA at 1 mg/kg (siRNA equivalents) on study days 1 and 8. The mice were euthanized on study day 22.

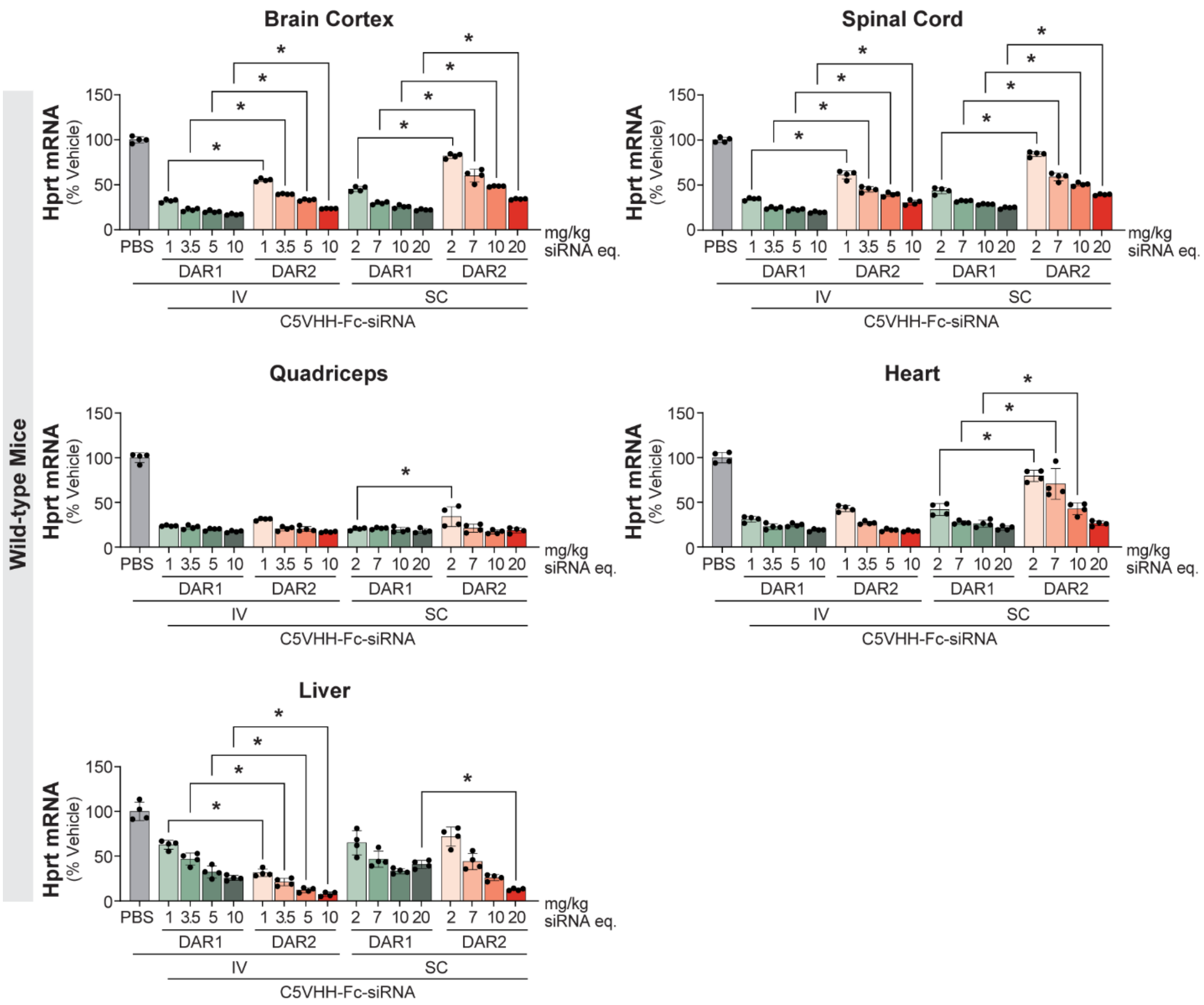


**Supplementary Figure 5:** DAR1 C5VHH-Fc-siRNA conjugate is more active in the CNS on a total siRNA mg/kg basis compared to DAR2 after systemic dosing in mice. Target Hprt mRNA levels measured by qRT-PCR in different tissues from wild-type mice dosed with DAR1 (one siRNA conjugated to C5VHH-Fc) or DAR2 (two siRNAs conjugated to C5VHH-Fc) conjugates, either twice (on study days 1 and 8) by intravenous (IV) injection at 1, 3.5 and 10 mg/kg siRNA equivalents (eq.), or once (on study day 1) via subcutaneous (SC) injection at 2, 7 and 20 mg/kg siRNA equivalents (i.e., keeping constant the total amount of compound dosed over the course of the experiment). The mice were euthanized, and tissues collected on study day 22 for both the IV and SC arms. DAR: drug to antibody ratio.

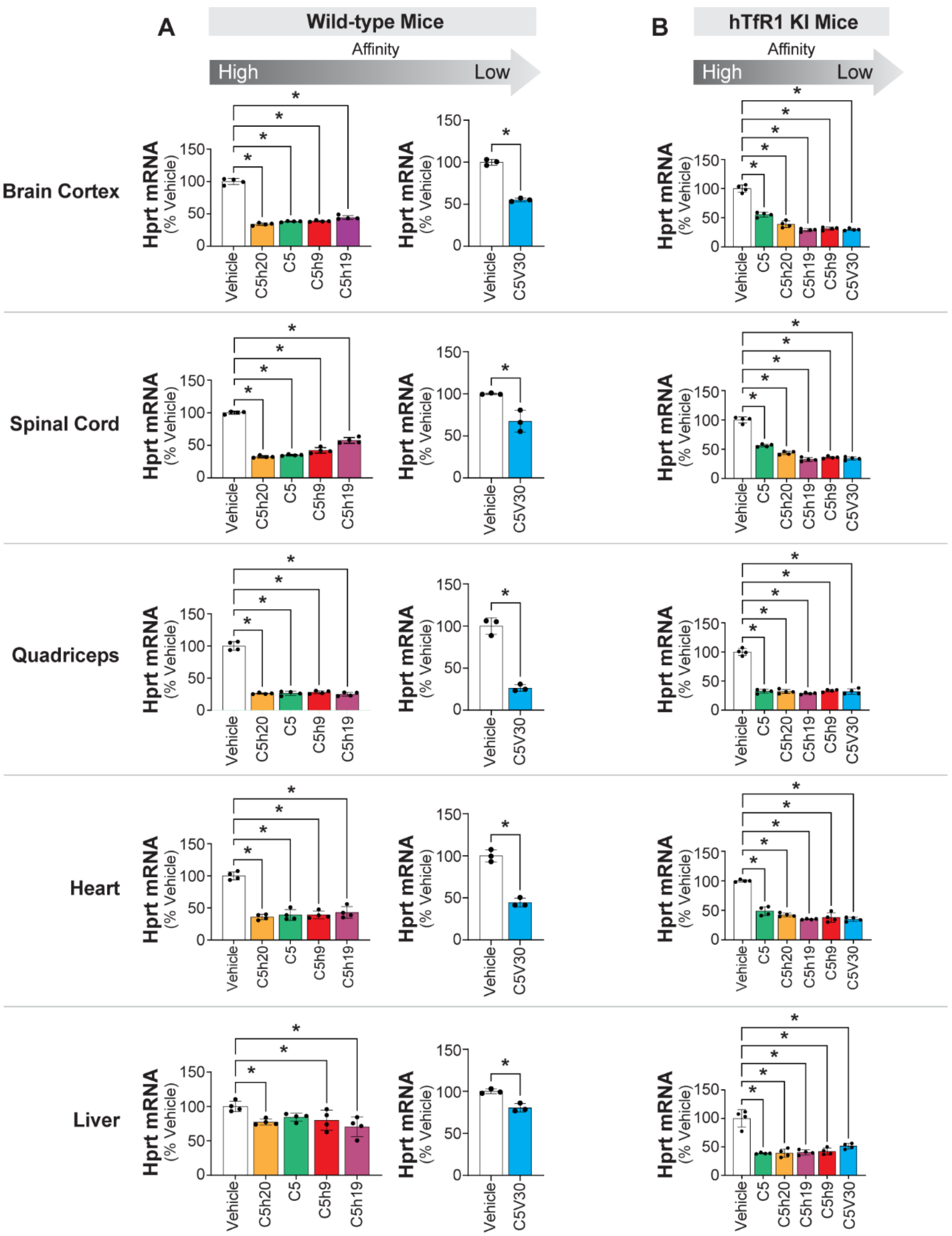


**Supplementary Figure 6:** The CNS activity of C5-series VHH-Fc-Hprt siRNA conjugates after IV dosing in wild-type and human TfR1 knock-in (hTfR1 KI) mice correlates with the binding affinity of the VHH clones for the cognate TfR1 receptor species. (**A**) Target Hprt mRNA levels measured by qRT-PCR in key

CNS and systemic tissues from wild-type mice dosed IV with different C5-series VHH-Fc-Hprt siRNA conjugates on study days 1 and 8 at 1 mg/kg (siRNA equivalents). The mice were euthanized on study day 22. As indicated by the arrow above the graphs, the conjugates are ordered based on the TfR1 binding affinity of the VHH variant for the mouse TfR1. For brain cortex, the same knockdown data reported here in a bar-graph representation were used to generate the plot in Figure 3A. (**B**) Target Hprt mRNA levels measured by qRT-PCR in key CNS and systemic tissues from hTfR1 KI mice. For brain cortex, the same knockdown data reported here in a bar-graph representation were used to generate the plot in Figure 3B.

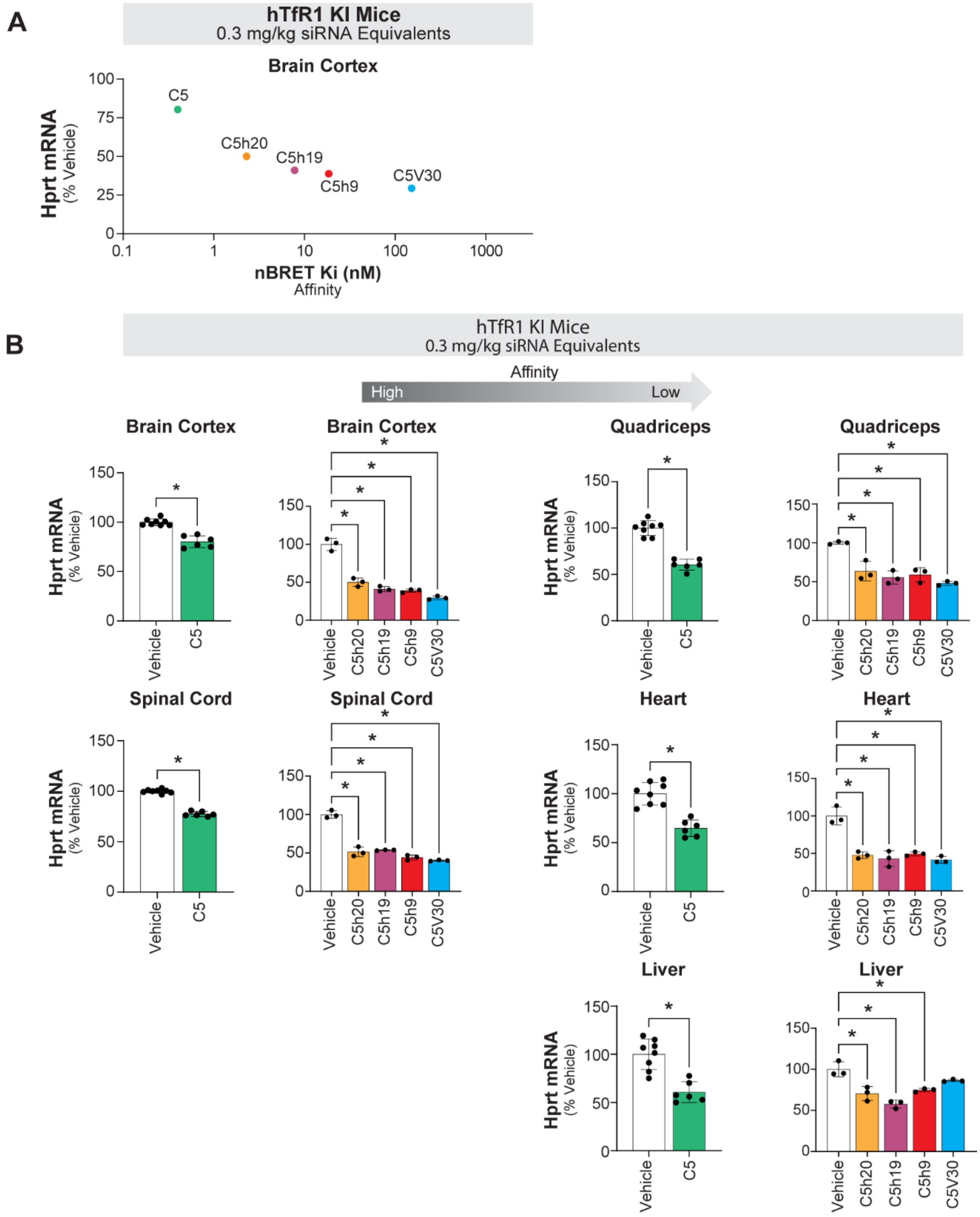


**Supplementary Figure 7:** Lower dose-level testing in human TfR1 knock-in mice helps differentiate the C5-family VHH clones based on CNS activity versus TfR1 binding affinity. (**A**) The target Hprt mRNA knockdown in brain cortex of human TfR1 KI (hTfR1 KI) homozygote mice (which express only human TfR1) was measured by qRT-PCR after IV dosing of various VHH-Fc-Hprt siRNA conjugates at 0.3 mg/kg (siRNA equivalents). The Hprt mRNA level is expressed as percentage of vehicle control animals and plotted versus the binding affinity (Ki) of each VHH-Fc-siRNA conjugate for the human TfR1, as measured by nBRET assay. The VHH clone is indicated, since it constitutes the part of the molecule that binds TfR1. (**B**) Target Hprt mRNA levels measured by qRT-PCR in key CNS and systemic tissues from hTfR1 KI mice dosed IV with the different C5-series VHH-Fc-Hprt siRNA conjugates on study days 1 and 8 at 0.3 mg/kg (siRNA equivalents). The mice were euthanized on study day 22. As indicated by the arrow above the graphs, the conjugates are ordered based on the TfR1 binding affinity of the VHH variant for the human TfR1. For brain cortex, the same knockdown data reported in B in a bar-graph representation were used to generate the plot in A.

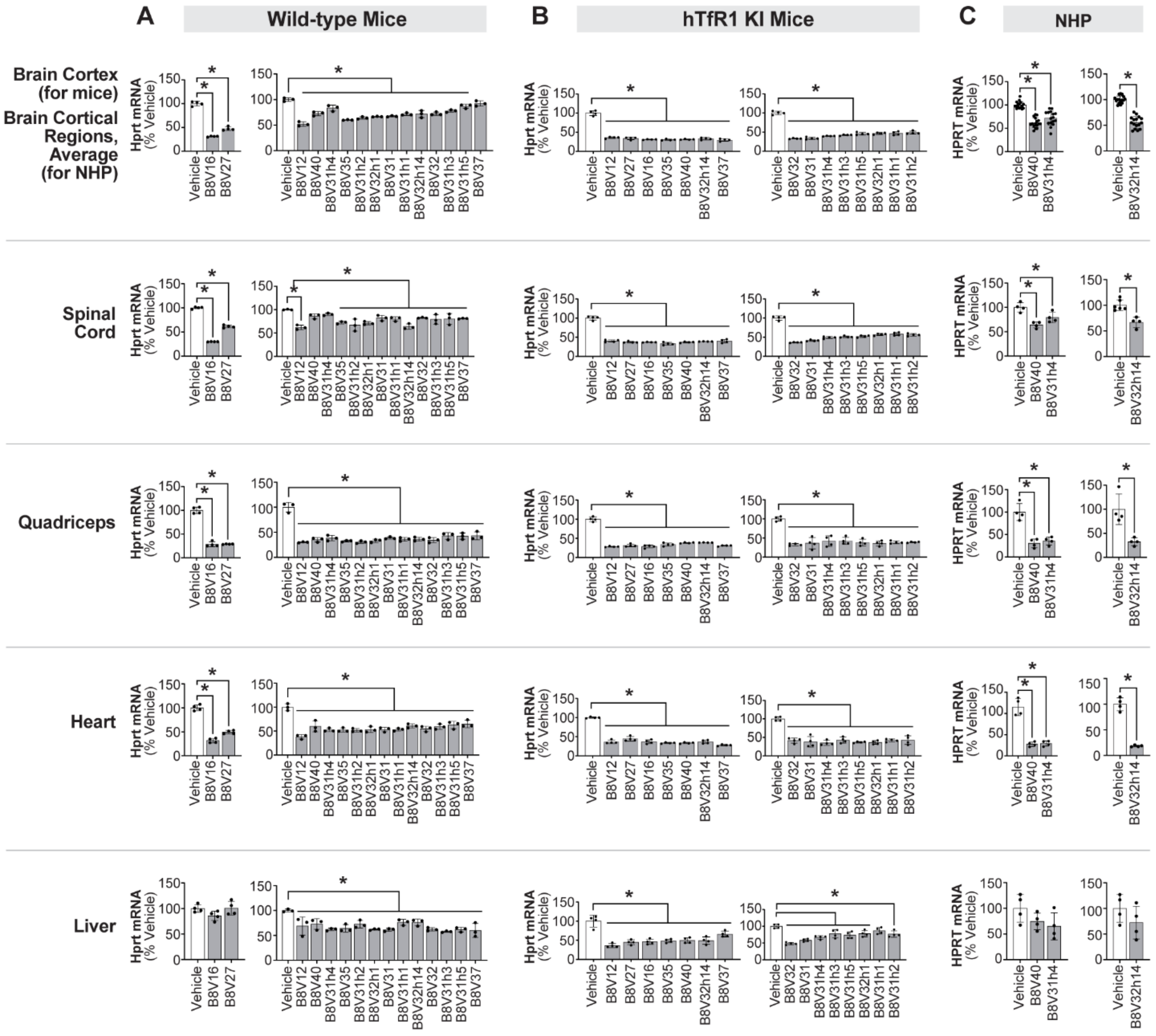


**Supplementary Figure 8:** TfR1-binding B8-series VHH-Fc-Hprt siRNA conjugates are active in the CNS after IV dosing in wild-type mice, human TfR1 knock-in mice, and NHP. (**A**) Target Hprt mRNA levels measured by qRT-PCR in key CNS and systemic tissues from wild-type mice dosed IV with different B8-series VHH-Fc-Hprt siRNA conjugates on study days 1 and 8 at 1 mg/kg (siRNA equivalents). The mice were euthanized on study day 22. Within each graph, the conjugates are ordered based on the TfR1 binding affinity of the VHH variant for the mouse TfR1. For brain cortex, the same knockdown data reported here in a bar-graph representation were used to generate the plot in Supplementary Figure 9A (for a sub-set of VHH clones). (**B**) Target Hprt mRNA levels measured by qRT-PCR in key CNS and systemic tissues from human TfR1 knock-in (hTfR1 KI) mice dosed IV with different B8-series VHH-Fc-Hprt siRNA conjugates on study days 1 and 8 at 1 mg/kg (siRNA equivalents). The mice were euthanized on study day 22. Within each graph, the conjugates are ordered based on the TfR1 binding affinity of the VHH variant for the human TfR1. For brain cortex, the same knockdown data reported here in a bar-graph representation were used to generate the plot in Supplementary Figure 9B (for a sub-set of VHH clones). (**C**) Target HPRT mRNA levels measured by qRT-PCR in key CNS and systemic tissues from NHPs dosed by IV infusion with different B8-series VHH-Fc-Hprt siRNA conjugates. The animals were dosed at 1 mg/kg (siRNA equivalents) on study days 1, 8, 15, and 22, and were euthanized on either day 24 (one animal) or day 36 (three animals). The two timepoints were combined due to the minimal differences in activity observed. For the graph on the left, the conjugates are ordered based on the TfR1 binding affinity of the VHH variant for the NHP TfR1. The top graph reports the average knockdown data from the combined brain cortical regions examined (frontal, motor, temporal, and occipital cortex). Each dot represents one cortical region in one animal (i.e., 4 dots for each animal, 16 dots total). For brain cortex, the same knockdown data reported here in a bar-graph representation were used to generate the plot in Supplementary Figure 9C.

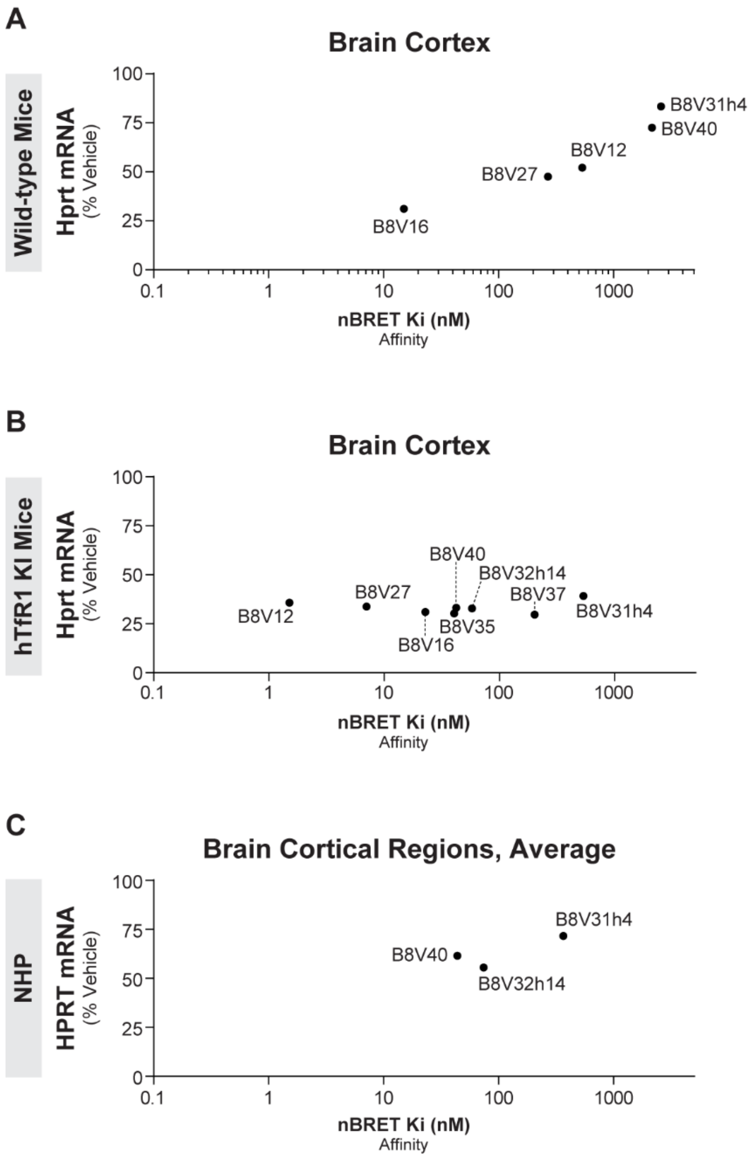


**Supplementary Figure 9:** The CNS activity of B8-series VHH-Fc-Hprt siRNA conjugates after IV dosing correlates with the binding affinity of the VHH clone for the cognate TfR1 receptor species. (**A**) The target Hprt mRNA knockdown in brain cortex of wild-type mice (which express only mouse TfR1) was measured by qRT-PCR after IV dosing of various B8-series VHH-Fc-Hprt siRNA conjugates at 1 mg/kg (siRNA equivalents). The Hprt mRNA level is expressed as percentage of vehicle control animals and plotted versus the binding affinity (Ki) of each VHH-Fc-siRNA conjugate for the mouse TfR1, as measured by nBRET assay. The VHH clone is indicated, since it constitutes the part of the molecule that binds TfR1. The same knockdown data plotted here are reported in a bar-graph representation in Supplementary Figure 8A. (**B**)

Similarly to A, the Hprt mRNA knockdown after IV dosing at 1 mg/kg in human TfR1 knock-in (hTfR1 KI) homozygote mice (which express only human TfR1) is plotted versus the binding affinity of the B8-series VHH conjugates for the human TfR1. The same knockdown data plotted here are reported in a bar-graph representation in Supplementary Figure 8B. (**C**) Target HPRT mRNA levels measured by qRT-PCR in brain cortical regions from NHPs after IV infusion with different B8-series VHH-Fc-Hprt siRNA conjugates. The animals were dosed at 1 mg/kg (siRNA equivalents) on study days 1, 8, 15, and 22, and were euthanized on either day 24 (one animal) or day 36 (three animals). The two timepoints were combined due to the minimal differences in activity observed. The average knockdown data from the combined brain cortical regions examined (frontal, motor, temporal, and occipital cortex) is reported. The same knockdown data plotted here are reported in a bar-graph representation in Supplementary Figure 8C.

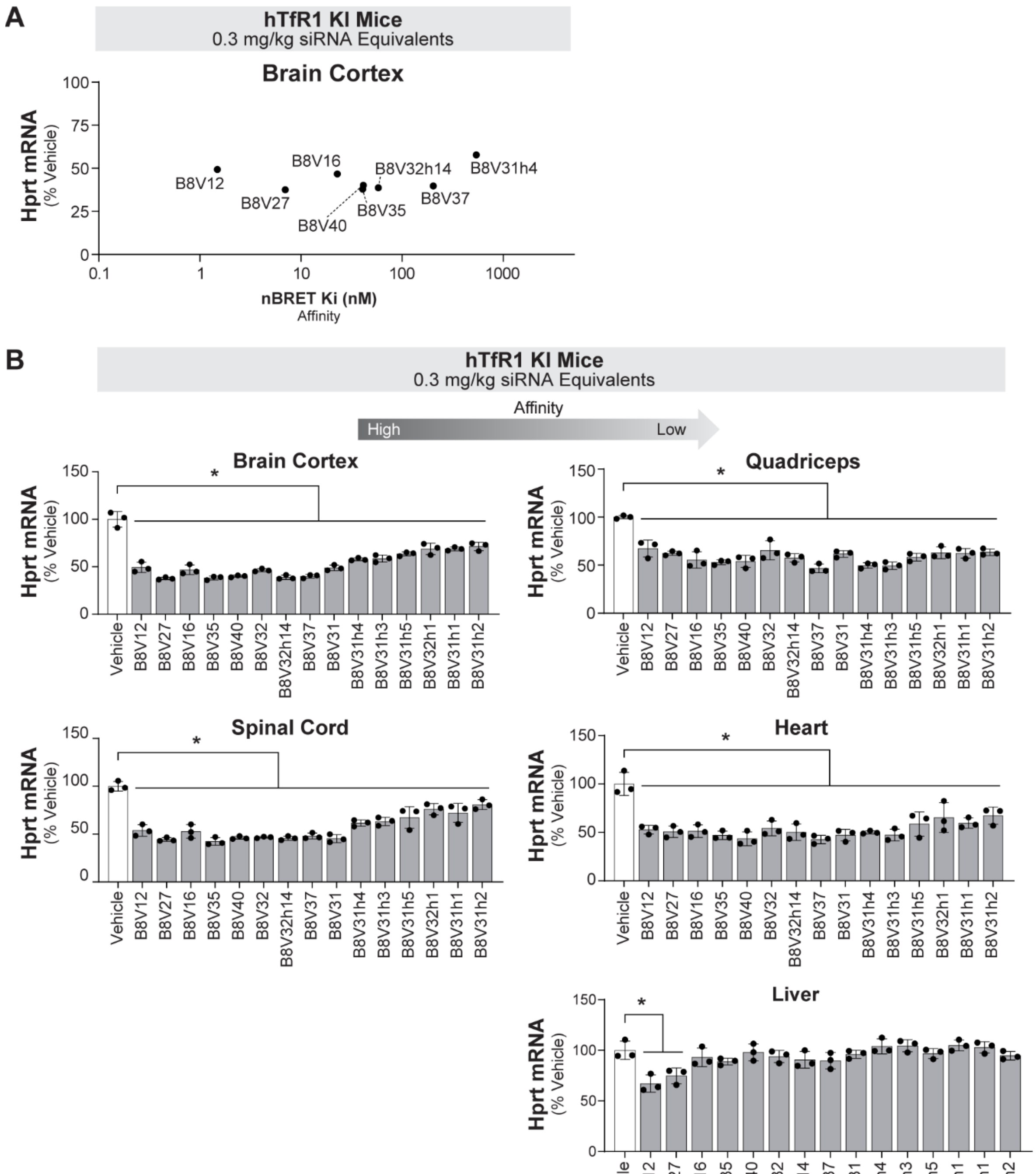


**Supplementary Figure 10:** Lower dose-level testing in human TfR1 knock-in mice helps differentiate the B8-family VHH clones based on CNS activity versus TfR1 binding affinity. (**A**) The target Hprt mRNA knockdown in brain cortex of human TfR1 knock-in (hTfR1 KI) homozygote mice (which express only human TfR1) was measured by qRT-PCR after IV dosing of various VHH-Fc-Hprt siRNA conjugates at

0.3 mg/kg (siRNA equivalents). The Hprt mRNA level is expressed as percentage of vehicle control animals and plotted versus the binding affinity (Ki) of each VHH-Fc-siRNA conjugate for the human TfR1, as measured by nBRET assay. The VHH clone is indicated, since it constitutes the part of the molecule that binds TfR1. (**B**) Target Hprt mRNA levels measured by qRT-PCR in key CNS and systemic tissues from hTfR1 KI mice dosed IV with the different B8-series VHH-Fc-Hprt siRNA conjugates on study days 1 and 8 at 0.3 mg/kg (siRNA equivalents). The mice were euthanized on study day 22. As indicated by the arrow above the graphs, the conjugates are ordered based on the TfR1 binding affinity of the VHH variant for the human TfR1. For brain cortex, the same knockdown data reported in B in a bar-graph representation were used to generate the plot in A.

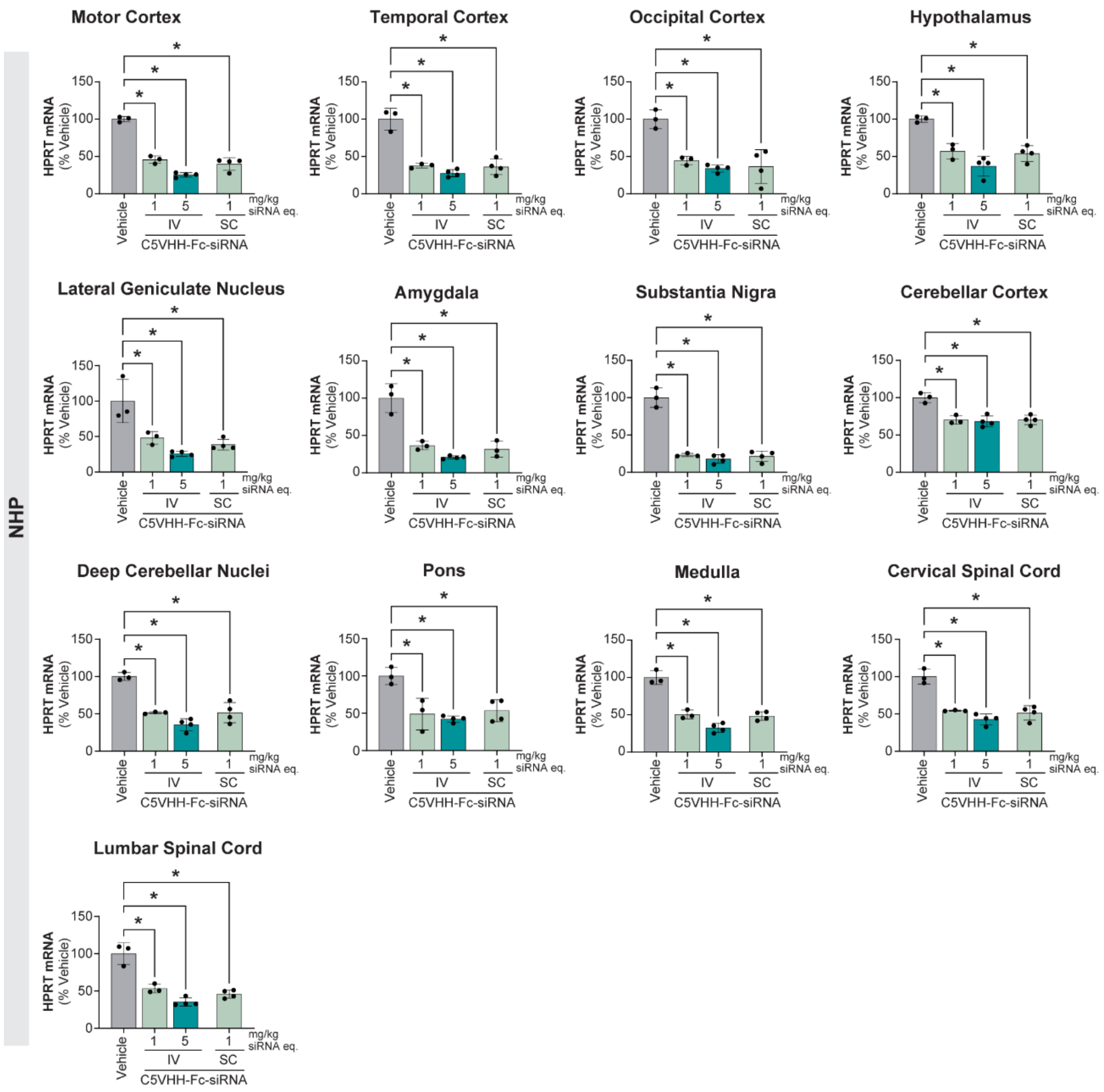


**Supplementary Figure 11:** IV and SC dosing of C5VHH-Fc-Hprt siRNA results in robust target mRNA knockdown in different brain regions and in spinal cord of NHPs. Target HPRT mRNA knockdown measured by qRT-PCR in various CNS tissues from NHPs dosed IV or SC with C5VHH-Fc-HPRT siRNA at the indicated dose levels, based on siRNA equivalents (eq.). The animals were dosed on study days 1, 8, 15, and 22, and they were euthanized on day 36. One animal in each of the two IV treatment groups was dosed as a sentinel, approximately 48 hours before the remaining animals in those groups and was euthanized on day 38. N equals 3 for the vehicle group and the 1 mg/kg IV group. N equals 4 for the 5 mg/kg IV group and the 1 mg/kg SC group. IV: dosing by intravenous infusion; SC: dosing by subcutaneous injection.

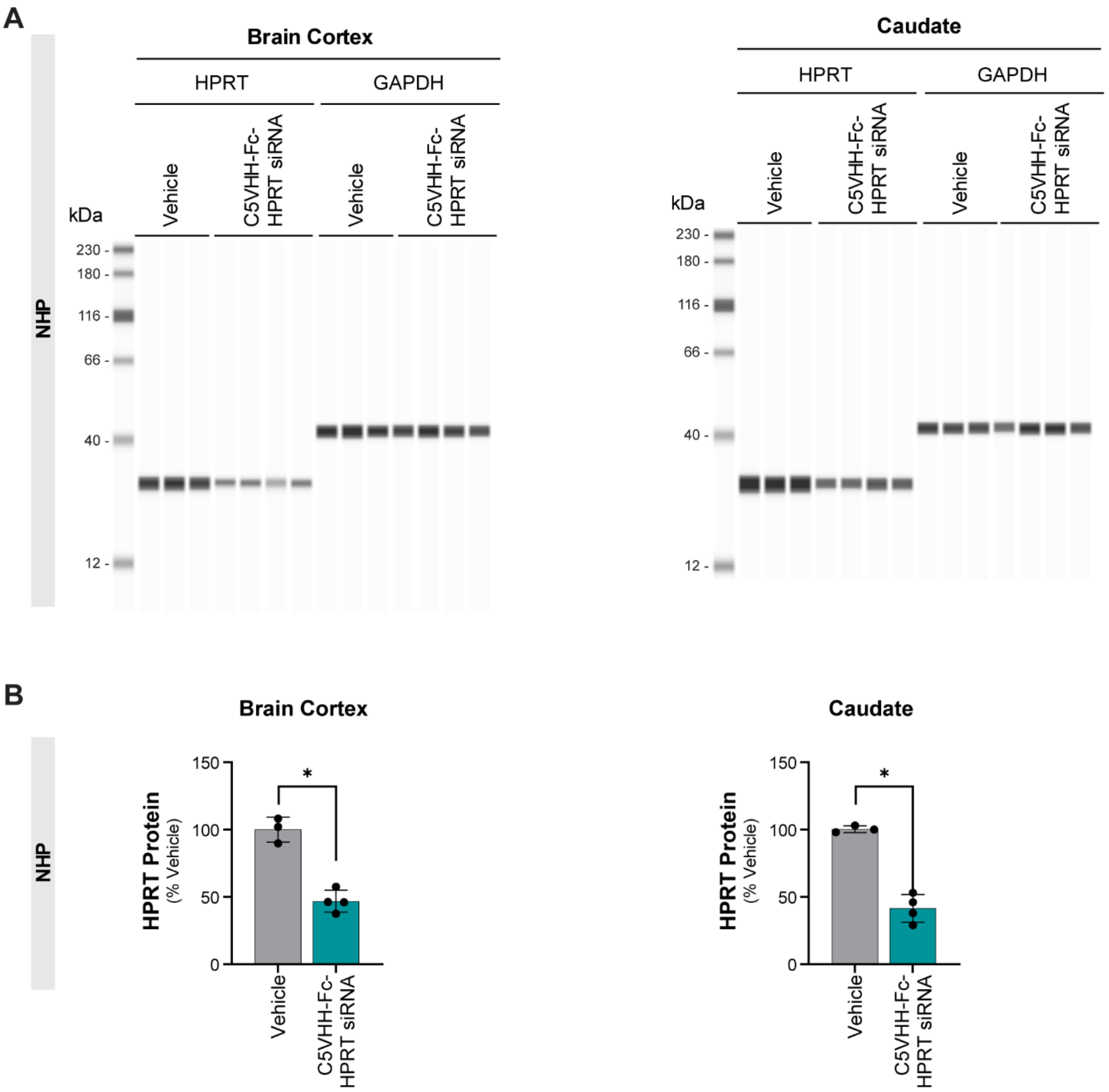


**Supplementary Figure 12:** IV dosed C5VHH-Fc-HPRT siRNA reduces the target HPRT protein in the CNS of NHPs. (**A**) HPRT protein levels were measured in brain cortex and caudate from NHPs dosed by intravenous infusion with C5VHH-Fc-HPRT siRNA at 5 mg/kg (siRNA equivalents) on study days 1, 8, 15, and 22. Animals were euthanized on day 36, and tissues were collected for analysis. HPRT protein was quantified by Western blot using the Simple Western automated Western blot system (Bio-Techne). (**B**) The intensity of the Western blot bands in A was analyzed using the integrated Simple Western software (Compass for Simple Western, Bio-Techne), and the HPRT signal was normalized to GAPDH as loading control. The HPRT protein levels were then expressed as percentage of vehicle-treated control animals.

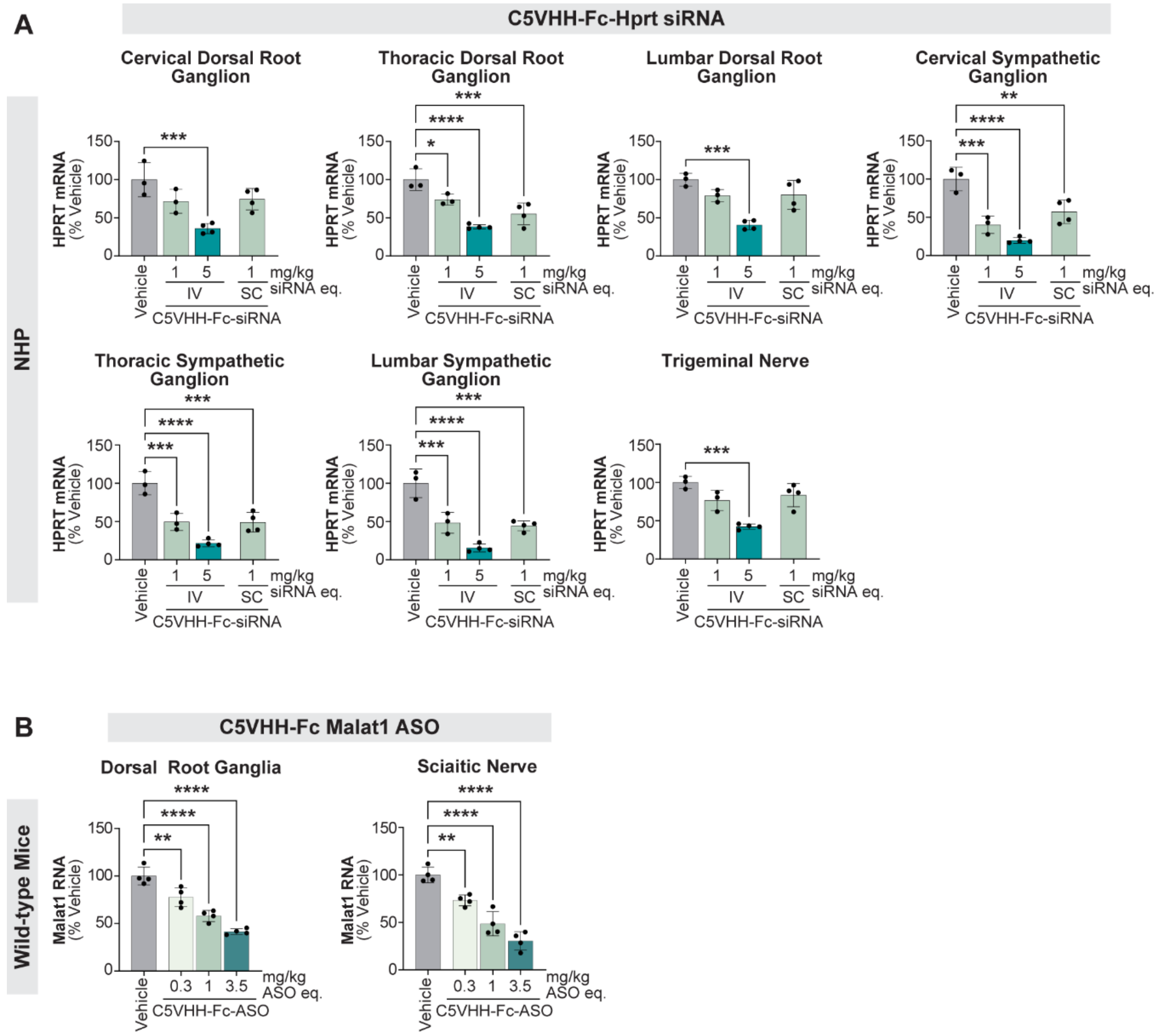


**Supplementary Figure 13:** C5VHH-Fc-HPRT siRNA is active in dorsal root ganglia and peripheral nerves of NHPs and wild-type mice after systemic dosing. (**A**) Target HPRT mRNA knockdown measured by qRT-PCR in dorsal root ganglia (DRG) and trigeminal nerve from NHPs dosed IV or SC with C5VHH-Fc-HPRT siRNA at the indicated dose levels, based on siRNA equivalents (eq.). The animals were dosed on study days 1, 8, 15, and 22, and were euthanized on day 36. N equals 3 for the vehicle group and the 1 mg/kg IV group. N equals 4 for the 5 mg/kg IV group and the 1 mg/kg SC group. IV: dosing by intravenous infusion; SC: dosing by subcutaneous injection. (**B**) Target Malat1 RNA knockdown measured by qRT-PCR in DRG and sciatic nerve from wild-type mice dosed IV with C5VHH-Fc-Malat1ASO (3-10-3 cEt gapmer) at the indicated dose levels, based on ASO equivalents (eq.). The mice were dosed on study days 1 and 8, and were euthanized on day 16.

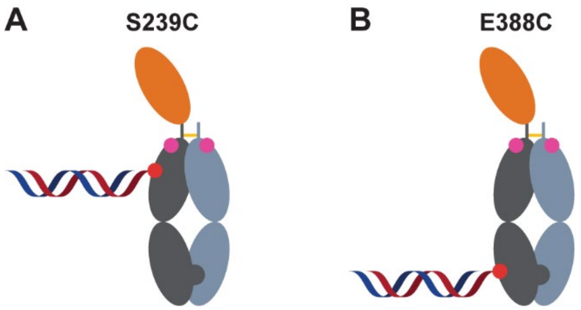


**Supplementary Figure 14:** Schematic representation of C5VHH–Fc-siRNA conjugates with different conjugation sites and strategies. (**A-B**) Illustrations of monovalent C5VHH–Fc constructs containing engineered cysteine residues at (**A**) S239C or (**B**) E388C, conjugated to an siRNA with a 5′ maleimide modification on the sense strand. The "knobs-into-holes" heterodimeric configuration is shown with the VHH ligand (orange) fused to the Fc (grey). T366W-bearing "knob" chains are dark grey while T366S, L368A, Y407V-bearing "hole" chains are light grey. Immune effector function modulating modifications are represented by a pink dot ("LALAPG": L234A, L235A, P329G). The conjugation site is represented by a red dot. Hinge disulfide bonds are represented by gold bars. The siRNA is shown as duplexed stands.

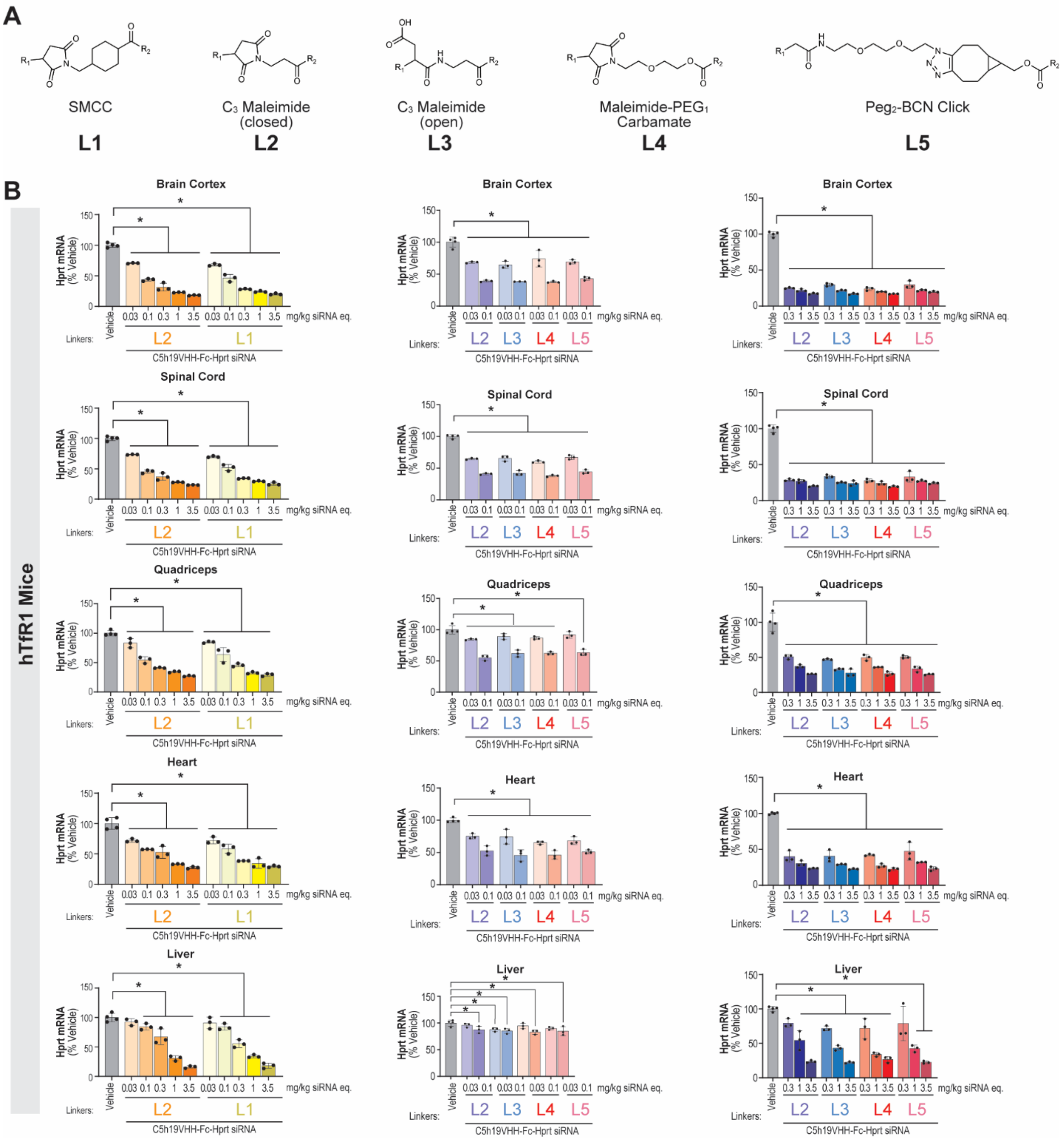


**Supplementary Figure 15:** Linker chemistry modifications do not improve CNS activity of systemically dosed VHH-Fc-Hprt siRNA conjugate. (**A**) Schematic representation of different linker chemistries. For each linker, R1 represents S from the engineered Cys239 of the VHH-Fc, and R2 represents the 5' NH from the hexylamino-siRNA. (**B**) Target Hprt mRNA levels measured by qRT-PCR in key CNS and systemic tissues from human TfR1 knock-in (hTfR1 KI) mice dosed with C5h19VHH-Fc-Hprt siRNA conjugates, where the siRNA is connected to the VHH-Fc at the S239C site using the different linker chemistries (L1-L5) shown in A. The mice were dosed IV on study days 1 and 8 at 0.03, 0.1, 0.3, 1.0, and 3.5 mg/kg siRNA equivalents (eq.) across three separate experiments (one graph per experiment). 0.5 mg of anti-mouse CD4 antibody were administered via intraperitoneal injection on study days 0 and 7. L2, which represents $C_3$ maleimide open ring, can have either a phosphodiester (orange bars) or a phosphorothioate (lavender bars) linkage connecting the siRNA to the C5h19VHH-Fc.

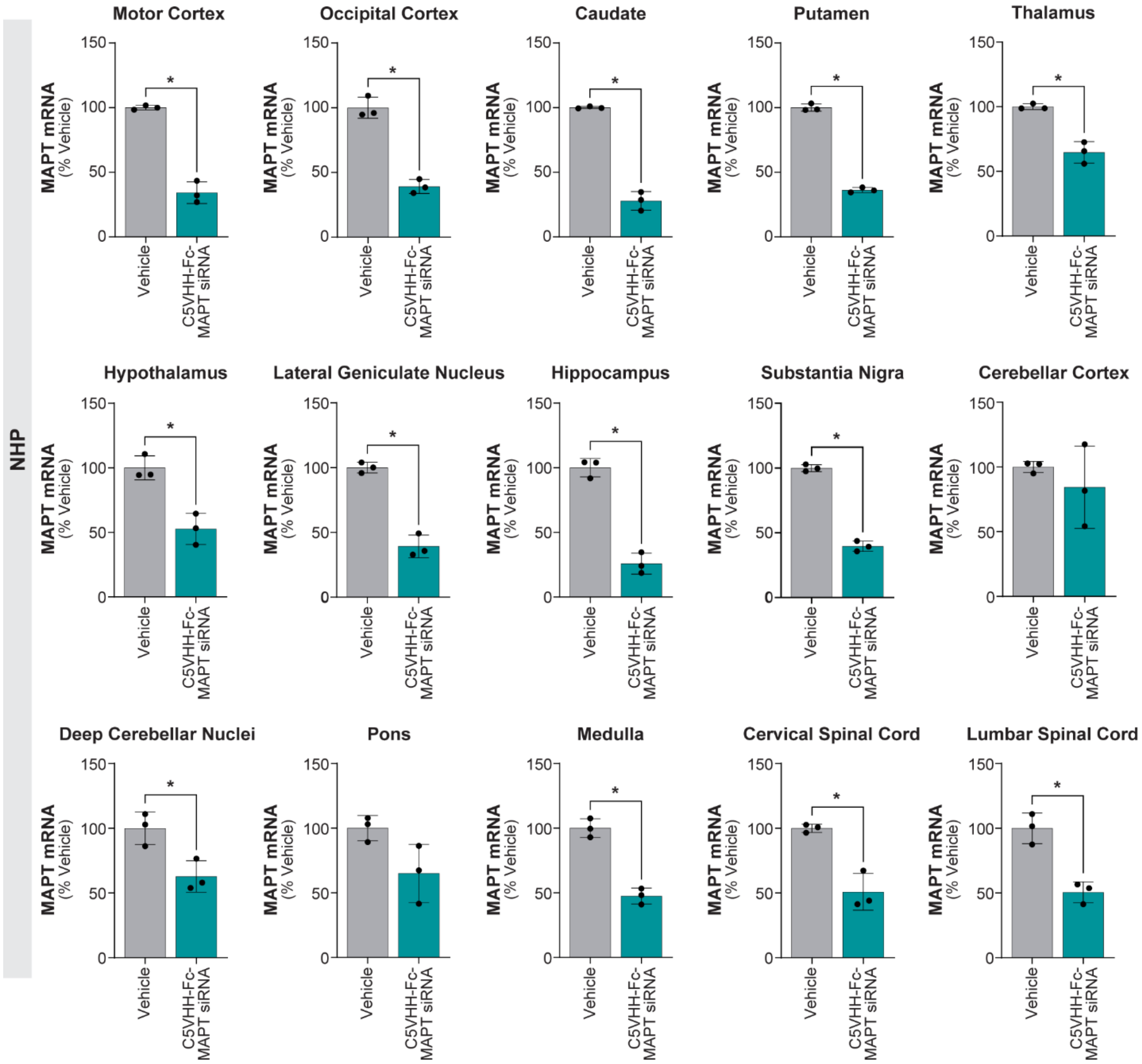


**Supplementary Figure 16:** IV dosed C5VHH-Fc-MAPT siRNA reduces the target MAPT mRNA in multiple brain regions and in spinal cord of NHPs. Target MAPT mRNA levels measured by qRT-PCR in different tissues from NHPs after IV infusion with C5VHH-Fc-MAPT siRNA at 5 mg/kg (siRNA equivalents). Levels are expressed as percentage of vehicle control animals. The NHPs were dosed on study days 1, 8, 15, and 22, and they were euthanized on day 36.

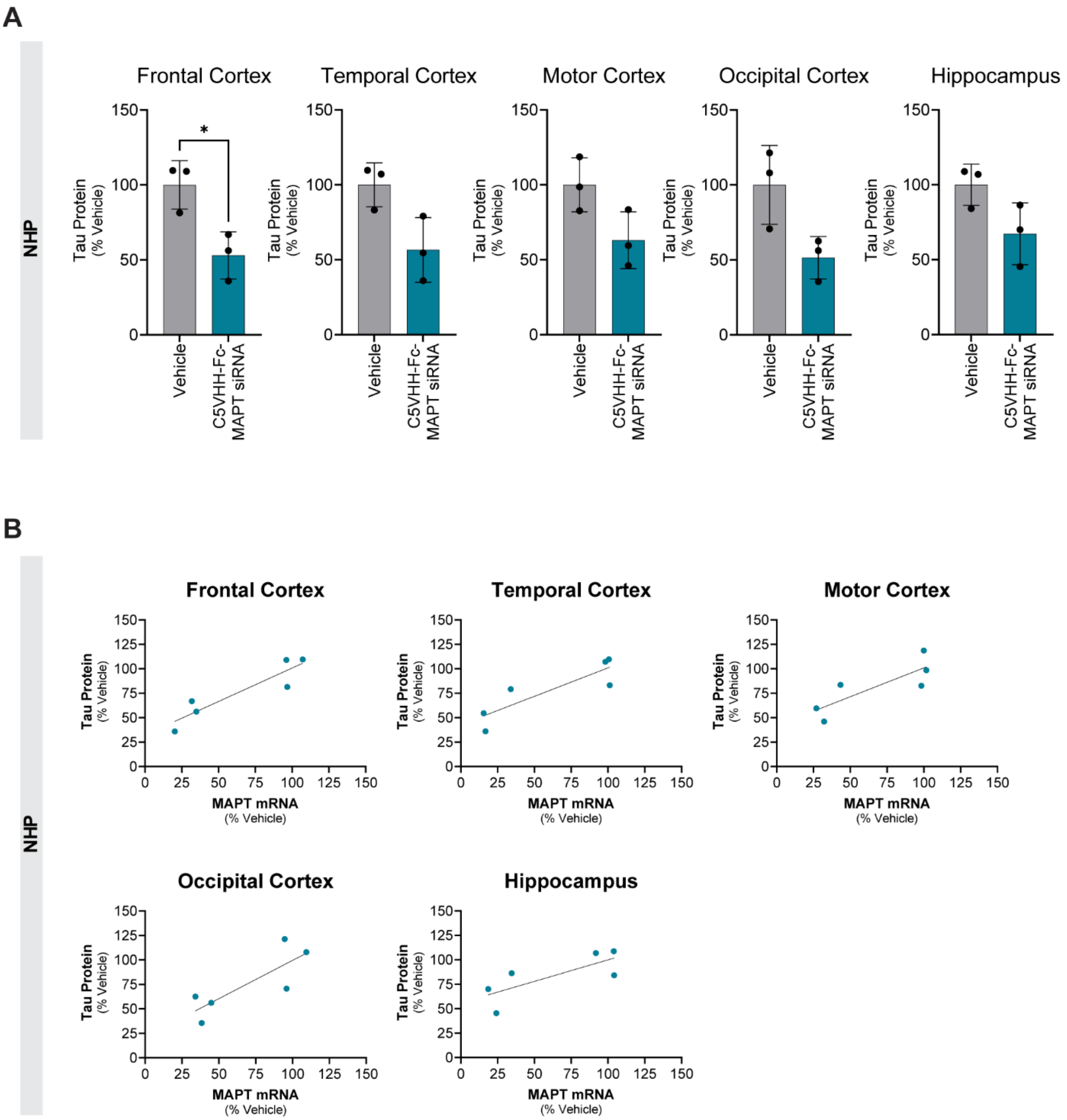


**Supplementary Figure 17:** IV dosed C5VHH-Fc-MAPT siRNA reduces target tau protein in the NHP brain. (**A**) Tau protein levels measured by ELISA in different brain regions from NHPs dosed via IV infusion with C5VHH-Fc-MAPT siRNA at 5 mg/kg (siRNA equivalents) on study days 1, 8, 15, 22. The animals were euthanized on day 36. N equals 3 per group. The tau protein levels are expressed as percentage of vehicle-treated NHPs. (**B**) Correlation plots of MAPT mRNA and tau protein levels across brain regions from NHPs treated by IV infusion with C5VHH-Fc-MAPT siRNA at 5 mg/kg siRNA equivalents. MAPT mRNA levels are plotted on the x-axis and tau protein levels on the y-axis. Values are expressed as percentages of vehicle-treated controls. The MAPT mRNA data are the same as those reported in Figure 6 and Supplementary Figure 16.

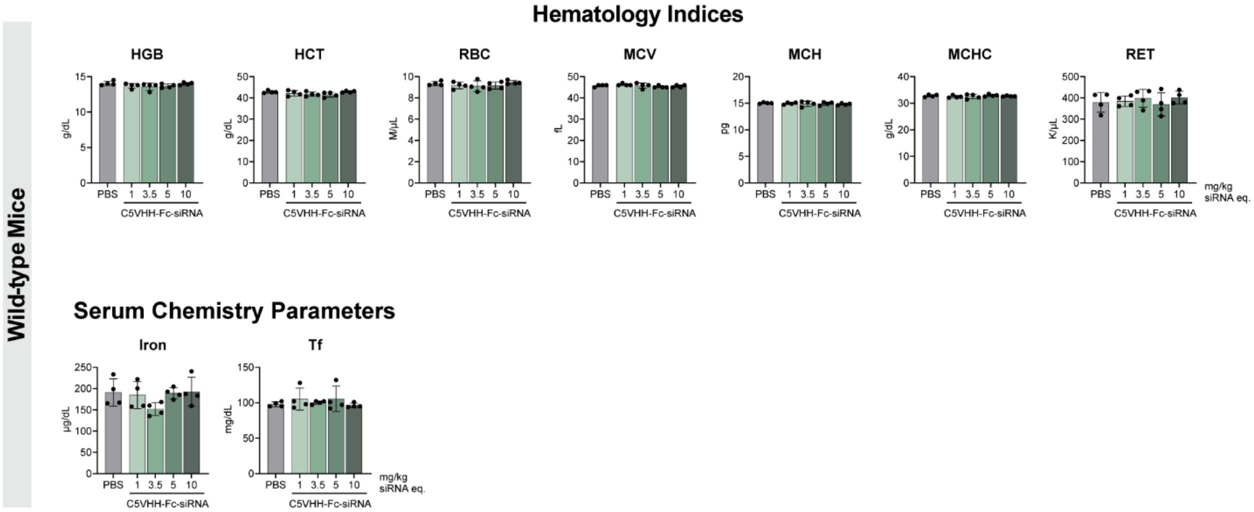


**Supplementary Figure 18:** Systemically dosed C5VHH-Fc-Hprt siRNA does not interfere with transferrin signaling, nor causes anemia in mice. Hematology indices and serum chemistry parameters from wild-type mice dosed with either vehicle (PBS) or C5VHH-Fc-Hprt siRNA at 1, 3.5, 5, or 10 mg/kg siRNA equivalents (eq.) via IV injection. HGB: hemoglobin; HCT: hematocrit; RBC: red blood cell count; MCV: mean corpuscular volume; MCH: mean corpuscular hemoglobin; MCHC: mean corpuscular hemoglobin concentration; RET: absolute reticulocyte count; Tf: transferrin.
